# Propofol induces a metabolic switch to glycolysis and cell death in a mitochondrial electron transport chain-dependent manner

**DOI:** 10.1101/181933

**Authors:** Chisato Sumi, Akihisa Okamoto, Hiromasa Tanaka, Kenichiro Nishi, Munenori Kusunoki, Tomohiro Shoji, Takeo Uba, Yoshiyuki Matsuo, Takehiko Adachi, Jun-Ichi Hayashi, Keizo Takenaga, Kiichi Hirota

## Abstract

The intravenous anesthetic propofol (2,6-diisopropylphenol) has been used for the induction and maintenance of anesthesia in operating rooms and for sedation in intensive care units. Although there is no widely accepted definition of propofol infusion syndrome (PRIS), PRIS is defined as the development of metabolic acidosis, rhabdomyolysis, hyperkalemia, hepatomegaly, renal failure, arrhythmia, and progressive cardiac failure. In vitro evidence suggests that PRIS is related to the impaired mitochondrial function. There are indications that preexisting mitochondrial disorders predispose to PRIS. However, the precise molecular mechanisms, including mitochondrial defects and a metabolic conversion by propofol, are largely unknown as yet. To elucidate the underlying cellular and molecular mechanisms of PRIS, we investigated the effects of propofol on the cellular metabolic mode and cell death. We demonstrated that clinically relevant concentrations of propofol, used within a clinically relevant exposure time, suppressed the mitochondrial function, caused the generation of reactive oxygen species, and induced a metabolic switch, from oxidative phosphorylation to glycolysis, by targeting complexes I and III of mitochondria. The data also indicated that a predisposition to mitochondrial dysfunction, caused by a genetic mutation or pharmacological suppression of the electron transport chain by biguanides such as metformin and phenformin, promoted the cell death and caspase activation induced by propofol.

## Introduction

Since its introduction into clinical practice in 1986, propofol (2,6-diisopropylphenol) has been used for the induction and maintenance of anesthesia in operating rooms and for sedation in intensive care units ^1^. Although propofol is considered a safe agent for anesthesia and sedation, a rare but severe complication can occur, especially in patients receiving high doses of the anesthetic for prolonged periods. A number of clinical reports have indicated a serious side effect of propofol, propofol infusion syndrome (PRIS) ^2^. Since PRIS was first described in 1992 ^3^, the clinical awareness and research interest into this disorder have continued to grow. The medical literature now includes over 100 published case reports, case series, and reviews. Nonetheless, the exact incidence and etiology of PRIS remain unclear at present. Although there is no widely accepted definition of PRIS, PRIS is defined as the development of metabolic acidosis (lactic acidosis), rhabdomyolysis, hyperkalemia, hepatomegaly, renal failure, arrhythmia, and progressive cardiac failure ^2^. There is a strong association between PRIS and propofol infusion at doses greater than 4 mg/kg/h and an exposure longer than 48 h, although the precise molecular mechanisms of PRIS have not been elucidated. In vitro evidence suggests that PRIS is related to impaired mitochondrial function ^4^. There are indications that preexisting mitochondrial disorders predispose to PRIS ^4-6^.

Moreover, studies using isolated mitochondria have demonstrated that propofol showed an effect on the mitochondrial respiratory chain. A decrease in the transmembrane electrical potential (ΔΨ) has been reported in liver mitochondria isolated from rats incubated with propofol ^7^. An increase of the oxygen consumption rate (OCR) has suggested that propofol acts as an uncoupler in oxidative phosphorylation (OXPHOS) ^8^. It has also been indicated that incubation of isolated mitochondria from rats with high concentrations of propofol (100 to 400 μM) resulted in a strong inhibition of the activity of complex I and, to a lesser degree, of complexes II and III ^9^. Other reports have also demonstrated a reduction of complex IV activity in skeletal muscles, which led to a hypothesis that a propofol metabolite causes a disruption of the respiratory chain ^10^. However, the precise molecular mechanisms, including the relationship between mitochondrial defects and metabolic reprogramming in the pathophysiology of PRIS, are largely unknown as yet. To investigate the underlying cellular and molecular mechanisms of PRIS, we investigated the effects of propofol on cell life and death, oxygen metabolism, and mitochondrial function using cells of various origins, including transmitochondrial cybrid cells harboring a mitochondrial DNA defect and mutations. We demonstrated that clinically relevant concentrations of propofol, used within a clinically relevant exposure time, suppressed the mitochondrial function, caused the generation of reactive oxygen species (ROS), and induced the metabolic reprogramming ^11^, from OXPHOS to glycolysis, by targeting complexes I and III of mitochondria. The data also indicated that a predisposition to mitochondrial dysfunction, caused by a genetic mutation or pharmacological suppression of the electron transport chain (ETC) in mitochondria by biguanides such as metformin and phenformin, promoted the cell death and caspase activation induced by propofol.

## Results

### Propofol induced cell death and activation of caspases in a concentration‐ and time-dependent manner

To determine whether propofol induces the cell death, we examined the concentration‐ and time-response relationship between propofol and cell death. Neuronal SH-SY5Y cells were treated with the indicated concentrations of propofol and for the indicated times. Cells were stained with propidium iodide (PI) and recombinant fluorescein isothiocyanate (FITC)-conjugated annexin V, and the proportion of dead cells was evaluated by flow cytometry. Concentrations of propofol equal to or greater than 50 μM induced the cell death within 6 h (Figs. 1a and 1b). Interestingly, 25 μM propofol induced the cell death at a longer incubation, 12 h (Fig. 1c). Next, activities of caspase-9 and caspase-3/7 were evaluated. Propofol at concentrations equal to or greater than 50 μM activated caspase-9 (Fig. 2a). As in the case of cell death and caspase-9, caspase-3/7 was activated by the treatment with 50 μM or greater propofol concentrations within 6 h. Importantly, 25 μM propofol, which did not induce caspase-3/7 activation within 6 h (Fig. 2b), induced the activation at 12 h (Fig. 2c). Next, a release of lactate dehydrogenase (LDH) was investigated (Fig. 2d). Within 6 h, only propofol at a concentration of 150 μM increased the LDH release. On the other hand, during 12-h incubation, not only 150 μM but also 50 and 100 μM propofol statistically significantly increased the LDH release. Measurement of the mitochondrial membrane voltage (ΔΨm) showed that propofol at concentrations equal to or greater than 50 μM decreased ΔΨm within 6 h (Fig. 2e). In addition, at concentrations higher than 50 μM propofol suppressed the cell viability, measured by an MTS [3-(4,5-dimethylthiazol-2-yl)-5-(3-carboxymethoxyphenyl)-2-(4-sulfophenyl)-2*H*-tetrazolium] assay, within 6 h (Supplementary Fig. 1a). 2,4-Diisopropylphenol, commonly known as 2,4-propofol, is an isomeric form of propofol, which does not show a hypnotic effect. We tested the effects of 2,4-diisopropylphenol on caspase-3/7 activation in SH-SY5Y cells. Similar to propofol, 2,4-diisopropylphenol induced caspase-3/7 activation within 6 h (Supplementary Fig. 1b). Interestingly, 25 μM 2,4-diisopropylphenol could activate caspase-3/7 within 6 h, suggesting that 2,4-diisopropylphenol is more toxic than propofol. Finally, the effects of propofol were investigated in cells of different origins, including mouse myoblast C2C12 cells (Supplementary Fig. 1c), human cervical cancer HeLa cells (Supplementary Fig. 1d), and Lewis lung carcinoma P29 cells (Supplementary Fig. 1e). Similar to its effects on SH-SY5Y cells, 50 and 100 μM propofol but not 12.5 or 25 μM propofol induced caspase-3/7 activation in the other cell lines within 6 h.

**Figure 1.**
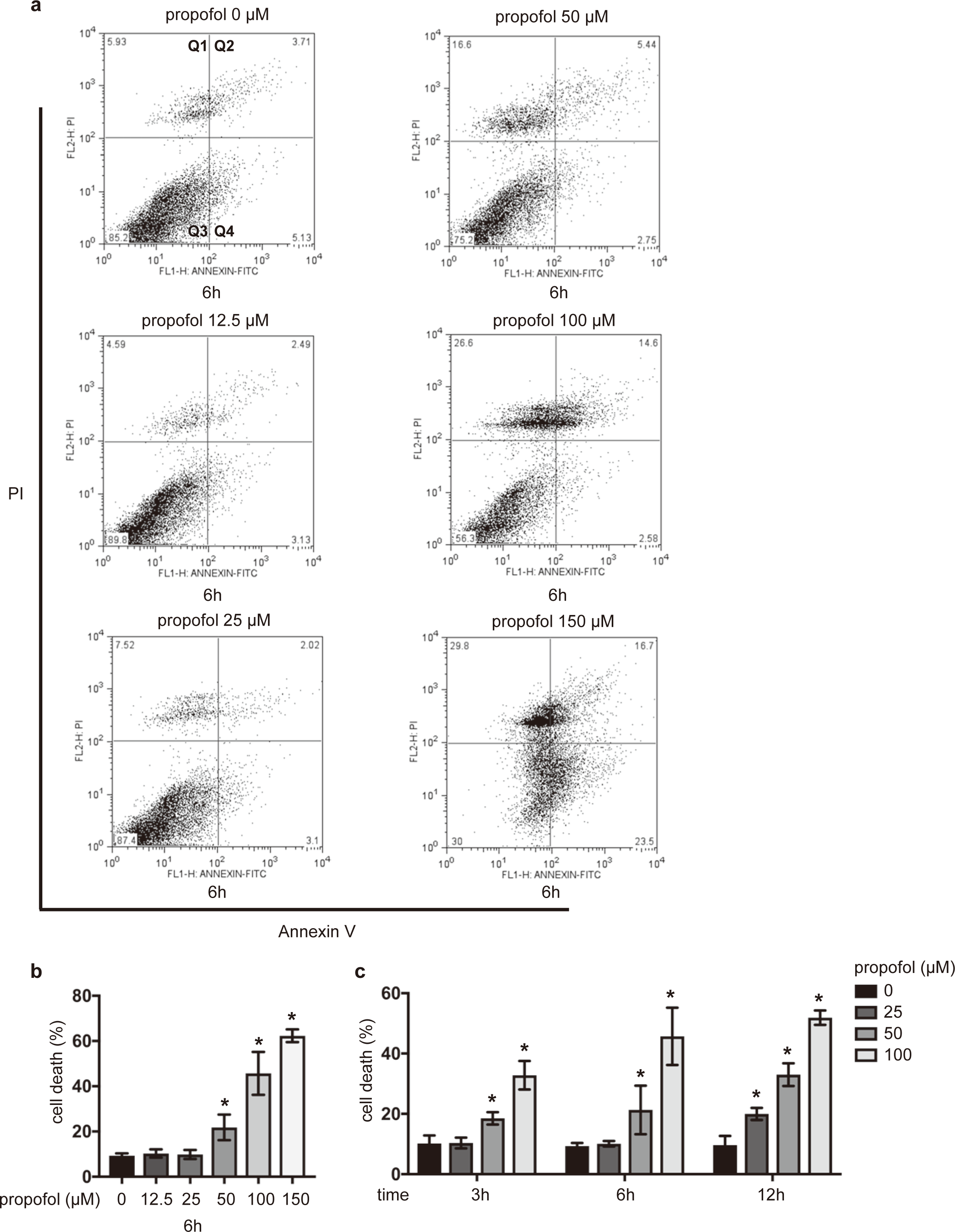
Propofol induced cell death in a concentration‐ and time-dependent manner. SH-SY5Y cells were exposed to the indicated concentrations (12.5, 25, 50, 100, and 150 μM) of propofol for 6 h (a and b) and 3, 6, and 12 h (c). Cells were harvested, and percentages of cell death were measured by flow cytometry. The ratio of PI-positive and/or annexin V-positive cells [(Q1 + Q2 + Q4)/(Q1 + Q2 + Q3 + Q4)] was used to calculate the percentage of dead cells (a and b) (n = 3). Data presented in (b and c) are expressed as the mean ± SD. Differences between treatment groups were evaluated by one-way ANOVA, followed by Dunnett’s multiple comparison test (b), or by two-way ANOVA, followed by Dunnett’s multiple comparison test (c). \**p* < 0.05 compared to the control cell population (incubation for 0 h, no treatment).

**Figure 2.**
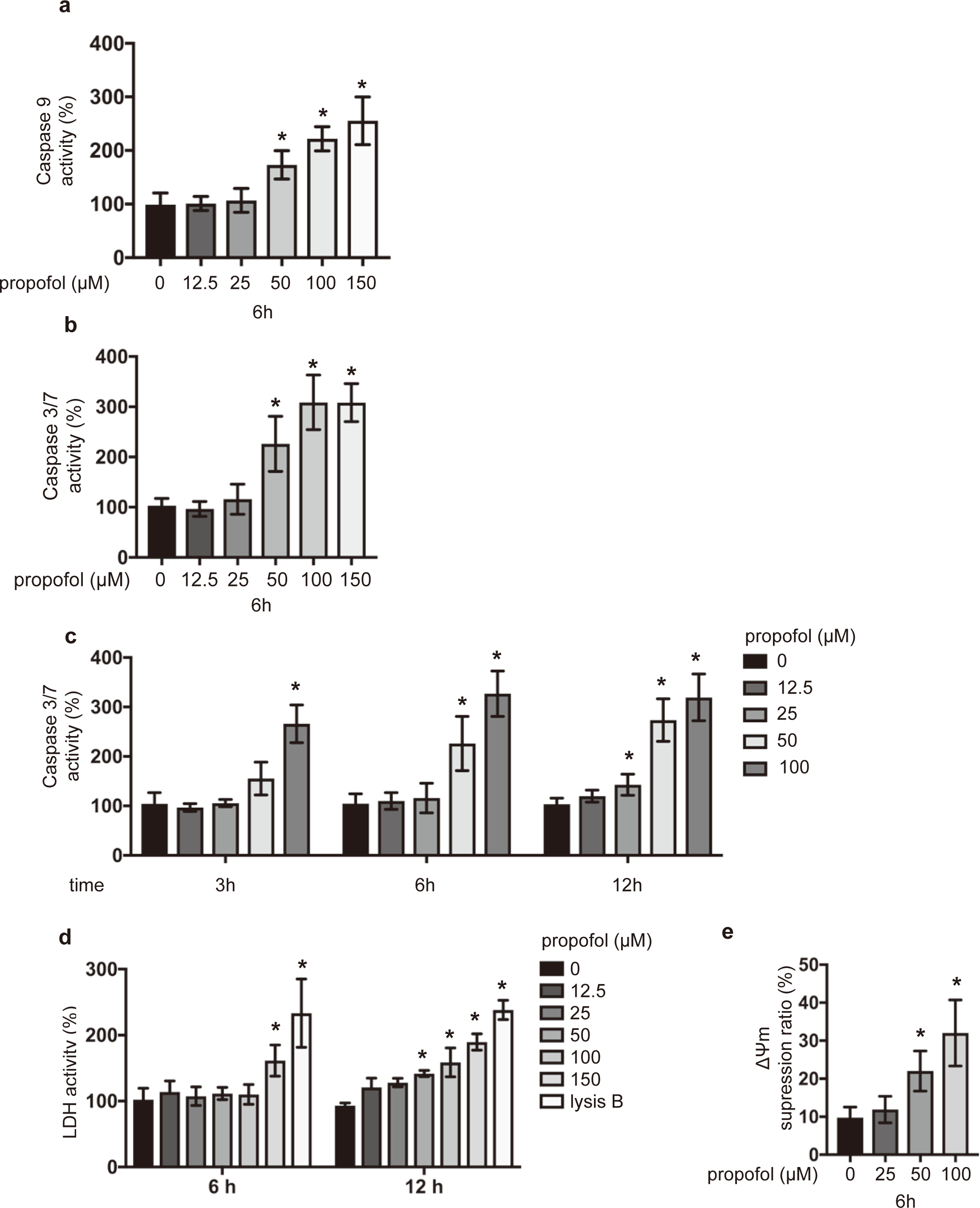
Propofol induced caspase activation in a concentration‐ and time-dependent manner. SH-SY5Y cells were exposed to the indicated concentrations (12.5, 25, 50, 100, and 150 μM) of propofol for 6 h (a and b) and 3, 6, and 12 h (c). Caspase-9 (n = 5) (a) and caspase-3/7 (n = 5) (b and c) activities in each treatment group at different time points.(d) SH-SY5Y cells were exposed to the indicated concentrations (12.5, 25, 50, 100, and 150 μM) of propofol for 6 and 12 h. The levels of LDH activity were assayed in culture supernatants (n = 3). Treatment with lysis buffer served as a control. (e) Average mitochondrial membrane potential (ΔΨm) of untreated cells and cells treated with the indicated concentrations (25, 50, and 100 μM) of propofol (n = 3) for 6 h. Values indicate the ratio [Q2/(Q2 + Q4)] of green JC-1 monomers (527 nm emission) to red aggregates (590 nm emission). Data presented in (a–g) are expressed as the mean ± SD. Differences between treatment groups were evaluated by one-way ANOVA, followed by Dunnett’s multiple comparison test (a, b, and e), or by two-way ANOVA, followed by Dunnett’s multiple comparison test (c and d). \**p* < 0.05 compared to the control cell population (incubation for 0 h, no treatment).

### Propofol suppressed oxygen metabolism and induced ROS generation

We investigated the effects of propofol on oxygen metabolism and glycolysis in SH-SY5Y cells by assaying OCR and the extracellular acidification rate (ECAR), which is a surrogate index for glycolysis. SH-SY5Y cells were preincubated with the indicated concentrations of propofol for the indicated periods. The mitochondrial OCR was significantly suppressed by the treatment with 50 μM propofol for 6 h (Figs. 3a and 3b, Supplementary Figs. 2a and 2c–f). Accordingly, the ECAR levels were significantly higher upon the treatment with 50 μM propofol (Figs. 3c and 3d, Supplementary Fig.2b). Propofol at a concentration of 25 μM, which exerted no significant effects within 6 h, suppressed OCR (Fig. 3e) and enhanced ECAR (Fig. 3f) after 12 h of incubation. Thus, the evidence indicates that propofol affects the oxygen metabolism in mitochondria. It has been reported that the disturbance of the mitochondrial ETC leads to the generation of ROS by cells ^12,13^. ROS generation was observed in SH-S5Y5 cells in response to propofol exposure within 3 and 6 h (Fig. 3g). Cell death was suppressed by the treatment with 10 mM *N*-acetylcysteine (Fig. 3h).

**Figure 3.**
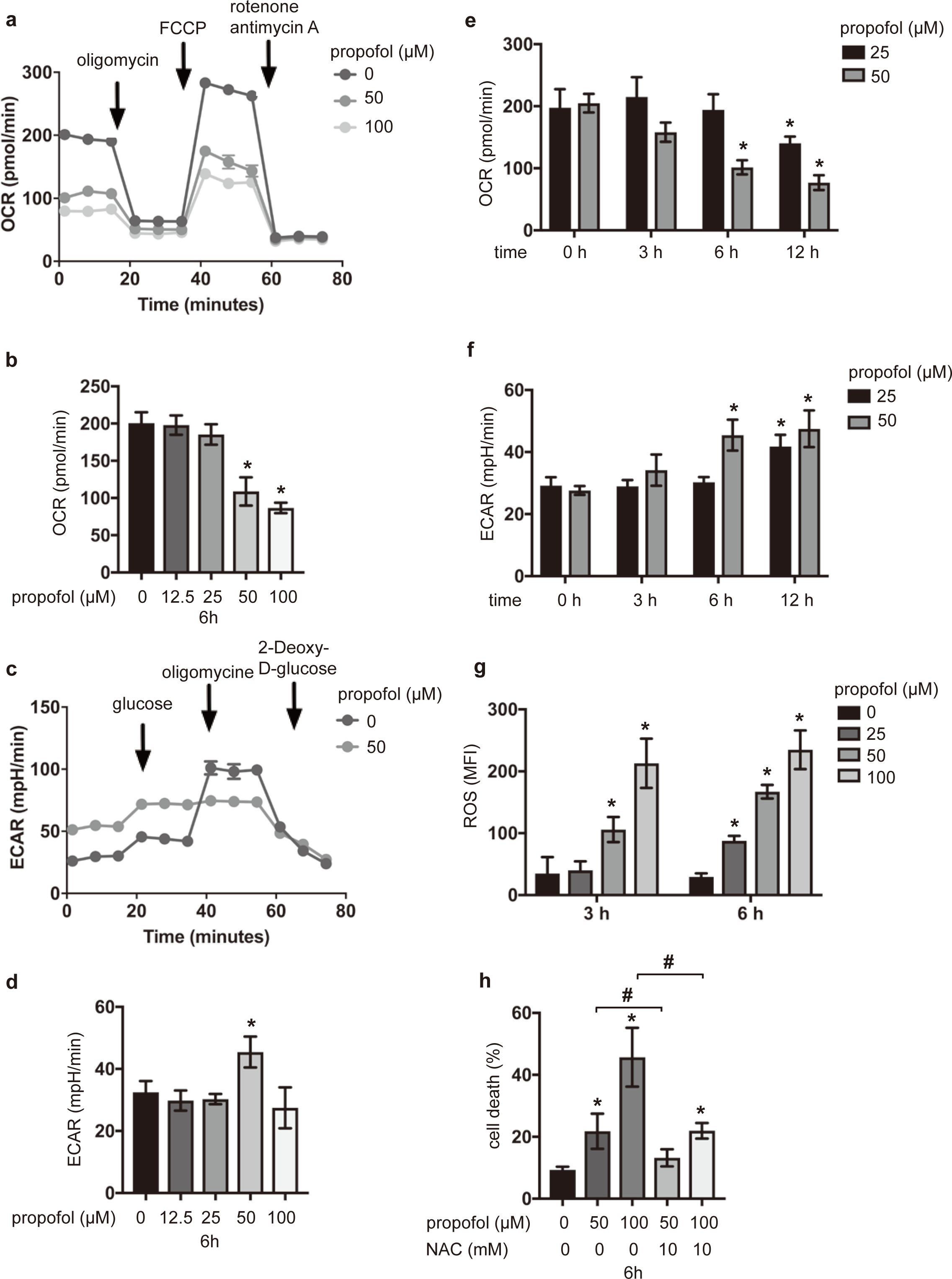
Oxygen metabolism and ROS generation in SH-SY5Y cells treated with propofol. OCR (a, b, and e) and ECAR (c, d, and f) in SH-SY5Y cells exposed to the indicated concentrations of propofol (12.5, 25, 50, and 100 μM) for 6 h (b and d) or 0, 3, 6, and 12 h (e and f). Data presented in (b, d–f) are expressed as the means ± SD. Differences between treatment groups were evaluated by one-way ANOVA, followed by Dunnett’s multiple comparison test (b and d), or by two-way ANOVA, followed by Dunnett’s multiple comparison test (e and f). (g) ROS production in SH-SY5Y cells exposed to 25, 50, and 100 μM propofol (n = 3) for 3 h. (h) SH-SY5Y cells were exposed to the indicated concentrations (50 and 100 μM) of propofol for 6 h with or without treatment with 10 mM *N*-acetylcysteine. Cells were harvested, and percentages of cell death were measured by flow cytometry. MFI: median fluorescence intensity; NAC: *N*-acetylcysteine. \**p* < 0.05 compared to the control cell population.

### Involvement of mitochondria in propofol-induced cell death and caspase activation

As shown in figure 3, propofol affected the mitochondrial ETC and intracellular oxygen metabolism in SH-SY5Y cells. To examine the involvement of mitochondria in propofol-induced cell death, we used the P29 cell line and its derivative ρ0P29, which lacks the mitochondrial DNA (mtDNA) ^14-16^. P29 and ρ0P29 cells were exposed to the indicated concentrations of propofol for 6 h, and then cell death (Fig. 4a) and the activity of caspase-3/7 (Fig. 4b) were assayed. Both cell death and caspase-3/7 assays indicated that, unlike P29 cells, ρ0P29 cells were completely resistant to 50 μM and partially resistant to 100 μM propofol after 6 h of incubation (Figs. 4a and 4b). Together with the results showing that propofol affects the ETC function of mitochondria, this evidence strongly suggests that mitochondria play a critical role and are one of the targets in propofol-induced cell death.

**Figure 4.**
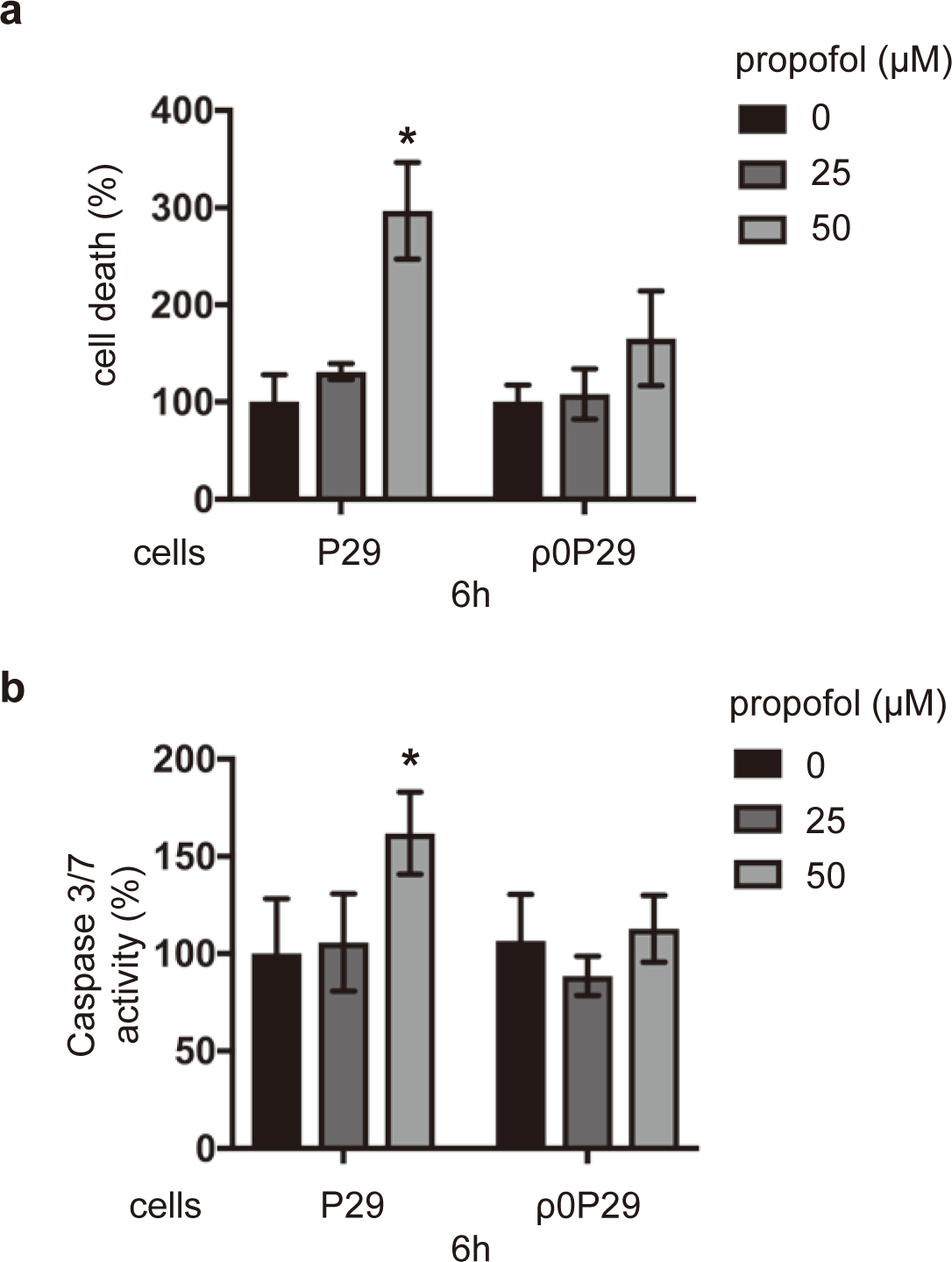
Involvement of functional mitochondria in propofol-induced caspase activation and cell death. P29 cells and cells of the ρ0P29 derivative lacking mtDNA were exposed to the indicated concentrations (25 and 50 μM) of propofol for 6 h. (a) Cells were harvested, and percentages of cell death were measured by flow cytometry. The ratio of PI-positive and/or annexin V-positive cells [(Q1 + Q2 + Q4)/(Q1 + Q2 + Q3 + Q4)] was used to calculate the percentage of dead cells (Supplementary Fig. 1a) (n = 3). (b) Caspase-3/7 activity in each treatment group (n = 3) at 6 h. Differences between treatment groups were evaluated by one-way ANOVA, followed by Dunnett’s multiple comparison test. \**p* < 0.05 compared to the control cell population; #*p* < 0.05 compared to the indicated experimental groups.

### Effects of propofol on the ETC complex-dependent OCR

Next, we examined the oxygen consumption, which depends on the activity of mitochondrial respiratory chain complexes I–IV in membrane-permeabilized and intact cells, using an Extracellular Flux Analyzer™ (Supplementary Fig. 3). OCR traces of mitochondrial respiration were detected using protocol A (Fig. 5a, Supplementary Fig. 3a) and protocol B (Fig. 5b, Supplementary Fig. 3b). The results indicated that propofol significantly suppressed the complex I‐ and complex III-dependent OCR but not the complex II‐ or complex IV-dependent OCR (Figs. 5c-5d).

**Figure 5.**
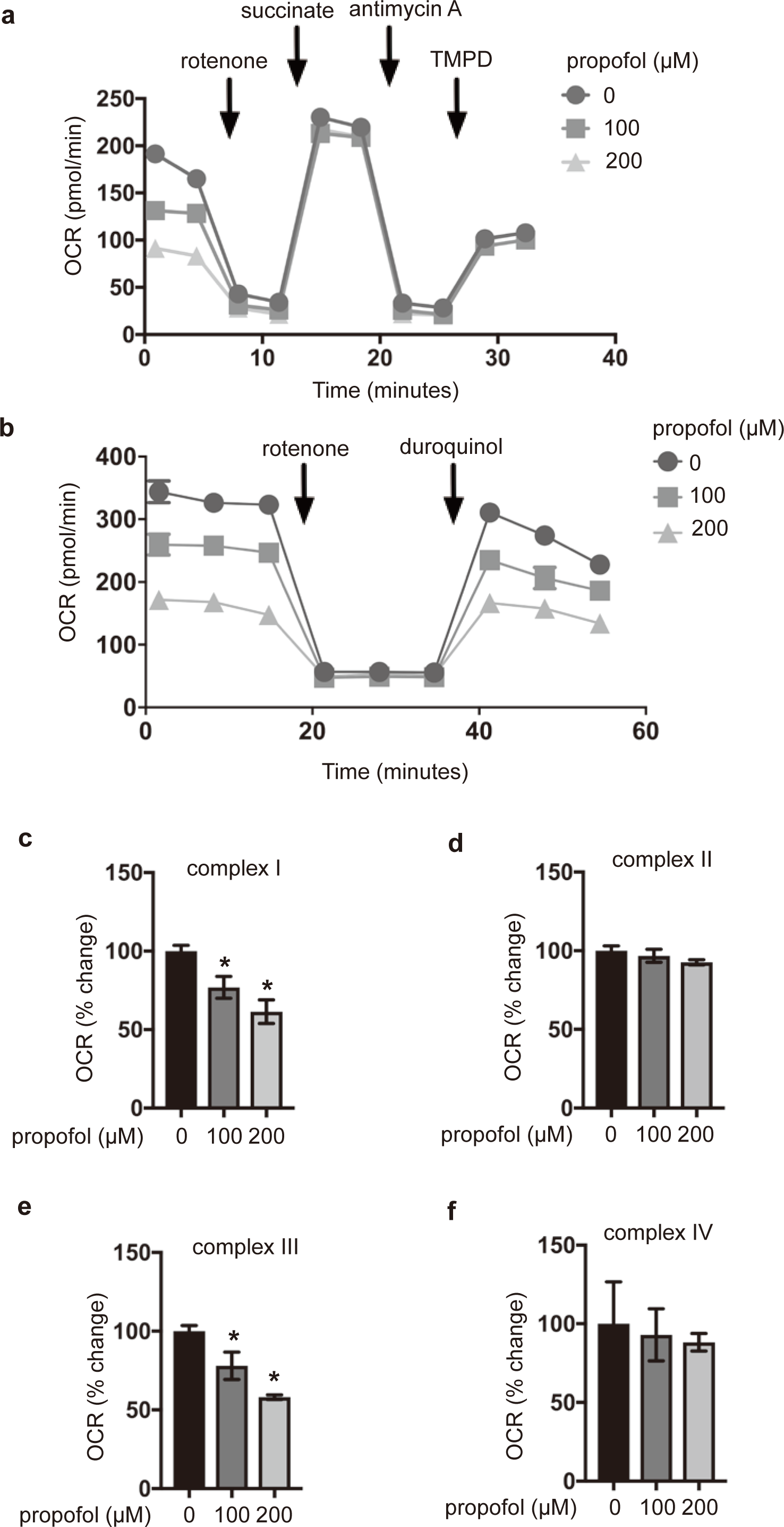
Effects of propofol on OCR driven by each complex of the mitochondrial electron transport chain. Representative OCR traces of mitochondrial respiration using protocol A (Supplementary Fig. 4a) and protocol B (Supplementary Fig. 4b). Mitochondrial ETC-mediated OCR, driven by complexes I (c), II (d), III (e), and IV (f), were assayed by a flux analyzer-based protocol. SH-SY5Y cells were exposed to 100 and 200 μM propofol for 6 h and subjected to the assay. Differences between treatment groups were evaluated by one-way ANOVA, followed by Dunnett’s multiple comparison test. \**p* < 0.05 compared to the control cell population.

### Mitochondrial ETC inhibitors synergistically enhanced propofol toxicity

We investigated the effects of several ETC inhibitors on the propofol-induced cell death in SH-SY5Y cells. Cells were exposed to rotenone (100 nM), antimycin A (25 μg/mL), or oligomycin (4 μM), with or without the indicated concentrations of propofol, for 6 h, and cell death was assayed by flow cytometry. Neither 100 nM rotenone, 25 μg/mL antimycin, 4 μM oligomycin, nor 12.5 μM or 25 μM propofol alone induced the cell death within 6 h (Fig. 6a). On the other hand, 12.5 and 25 μM propofol induced the cell death in the presence of rotenone, antimycin A, and oligomycin (Fig. 6a). Similarly, 12.5 and 25 μM propofol with rotenone, antimycin A, and oligomycin induced the caspase-3/7 activity within 6 h (Fig. 6b). The evidence indicates that cooperative inhibition of mitochondria by both propofol and the ETC inhibitors induces cell death at even clinically relevant concentrations of propofol within 6 h.

**Figure 6.**
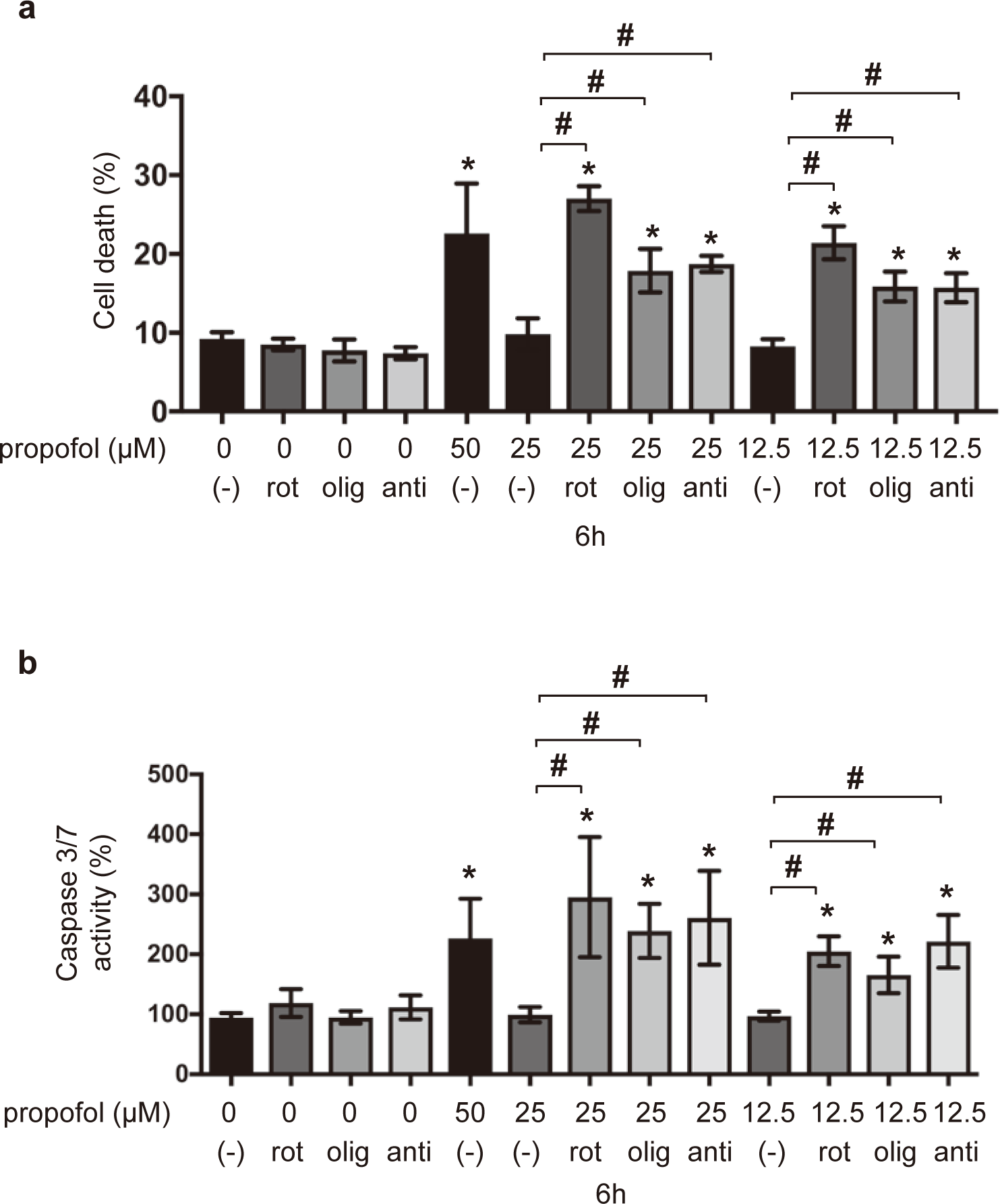
Synergistic effects of propofol and mitochondrial ETC inhibitors on caspase activity and cell death. Levels of caspase-3/7 activity and cell death of SH-SY5Y cells treated with propofol and mitochondrial ETC inhibitors. Cells were treated with 12.5, 25, or 50 μM propofol and with either 100 nM rotenone, 4 μM oligomycin, or 25 μg/mL antimycin A and subjected to (a) a cell death assay and (b) a caspase-3/7 activity assay (n = 3).Percentages of cell death were measured by flow cytometry. The ratio of PI-positive and/or annexin V-positive cells [(Q1 + Q2 + Q4)/(Q1 + Q2 + Q3 + Q4)] was used to calculate the percentage of dead cells (Supplementary Fig. 1a) (n = 3). All data are expressed as the means ± SD. \**p* < 0.05 compared with control cells (no treatment); #p < 0.05 compared with the indicated groups. rot: rotenone; olig: oligomycin; anti: antimycin A.

### Genetic predisposition to mitochondrial dysfunction increased propofol-induced caspase activation and cell death

For further study, we used transmitochondrial cybrids carrying mtDNA with defined pathogenic mutations. mtDNA donors and ρ0P29 cells were used to obtain transmitochondrial cybrids P29mtA11, P29mtB82M, P29mtCOIM, and P29mtΔ (Tables 2 and 3) ^16^, and oxygen metabolism profiles of these clones were examined (Figs. 7a and 7b). These transmitochondrial cybrids were exposed to the indicated concentrations of propofol for 6 h to investigate the cell death and caspase-3/7 activation. Interestingly, in contrast to parental P29 cells, the cell death flow cytometry assay indicated that 25 μM propofol induced the cell death within 6 h in P29mtA11, P29mtB82M, and P29mtCOIM cells (Fig. 7c). Caspase-3/7 activation was also induced in P29mtA11, P29mtB82M, and P29mtCOIM cells even with 25 μM propofol (Fig. 7d). In contrast, P29mtΔ cells, similar to ρ0P29 cells, were completely resistant to 50 μM propofol and partially resistant to 100 μM propofol (Fig. 7d). Thus, our experimental results indicated that cells harboring genetic mutations in mtDNA (*ND6* in complex I or *COI* in complex IV) were more susceptible to propofol.

**Figure 7.**
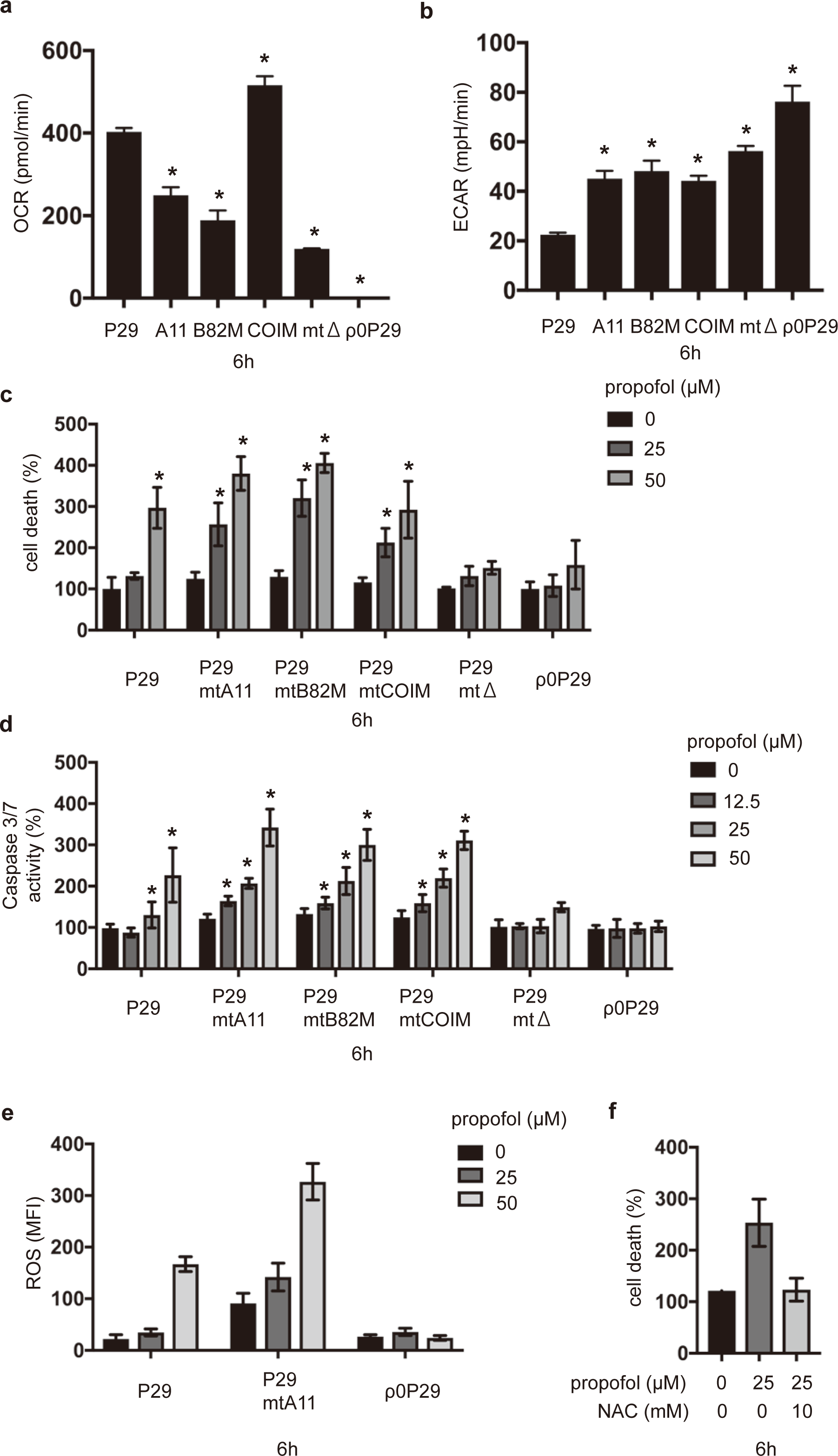
Effects of propofol on caspase activity and cell death in various transmitochondrial cybrid cells. (a) OCR and (b) ECAR of P29, its cybrid cells, and ρ0 cells. (c and d) P29, its cybrid cells, and ρ0 cells were exposed to the indicated concentrations (12.5, 25, and 50 μM) of propofol for 6 h. Cells were harvested, and percentages of cell death were measured by flow cytometry. The ratio of PI-positive and/or annexin V-positive cells [(Q1 + Q2 + Q4)/(Q1 + Q2 + Q3 + Q4)] was used to calculate the percentage of dead cells (n = 3) (c). Caspase-3/7 activity in each treatment group (n = 3) at 6 h (d). (e) P29, P29mtA11 and ρ0 cells were exposed to 25 and 50 μM propofol for 6 h and subjected to ROS assay. (f) P29mtA11 cells were exposed to 25 μM propofol with or without 10 mM NAC for 6 h. Cells were harvested, and percentages of cell death were measured by flow cytometry. Data presented in (a–f) are expressed as the means ± SD. Differences between treatment groups were evaluated by one-way ANOVA, followed by Dunnett’s multiple comparison test (a, b, e and f) or by two-way ANOVA, followed by Dunnett’s multiple comparison test (c and d). \**p* < 0.05 compared to the control cell population. A11: P29mtA11 cells; B82M: P29mtB82M cells; COIM: P29mtCOIM cells; mtΔ: P29mtΔ cells.

ROS generation was investigated in P29 cells, P29mtA11 cells and ρ0P29 cells in response to propofol exposure. Cells were exposed to 25 μM propofol for 6 h. P29mtA11 cells produced more ROS compared to P29 cells. 25 μM propofol, which did not produce ROS in P29 cells, induced generation of more ROS in P29mtA11 cells. In contrast, ρ0P29 cells did not produce ROS even under 50 μM propofol.

Accordingly, cell death induced by 25 μM propofol was suppressed by treatment with 10 mM NAC.

### Pharmacological suppression of mitochondrial ETC increased propofol-induced cell death and caspase activation

The biguanides metformin and phenformin are widely used to reduce high blood sugar levels caused by diabetes ^17^. Metformin and phenformin have also been shown to suppress complex I of ETC, which is used by cells to generate energy ^18-22^. To confirm the effect of blockade of ETC, SH-SY5Y cells were pretreated with 2.5–20 mM metformin or 5–15 μM phenformin for 6 h, with or without the indicated concentrations of propofol, and then the cells were tested using the OCR and ECAR assays. Incubation for 6 h with either 2.5 mM metformin or 5 μM phenformin did not affect OCR (Fig. 8a and Supplementary Fig. 4a) and ECAR (Fig. 8b and Supplementary Fig. 4b) in SH-SY5Y cells. In contrast, not only 25 μM but also 12.5 μM propofol, which did not affect OCR or ECAR alone, significantly decreased OCR (Fig. 8a) and increased ECAR (Fig. 8b) in the presence of 2.5 mM metformin. Next, we investigated ROS generation upon cell exposure to 25 μM propofol with or without 5 mM metformin. Exposure to 25 μM propofol did not induce ROS generation without metformin. With metformin, however, 25 μM propofol induced ROS generation (Fig. 8c). Then, SH-SY5Y cells were tested for cell death (Fig. 8d) and caspase 3/7 activation (Fig. 8e) after treatment with 5 mM metformin or 5 μM phenformin (Supplementary Figs. 4c and 4d) for 6 h, with or without the indicated concentrations of propofol. Metformin at 5 mM and phenformin at 5 μM increased the propofol-induced cell death and caspase-3/7 activation.

**Figure 8.**
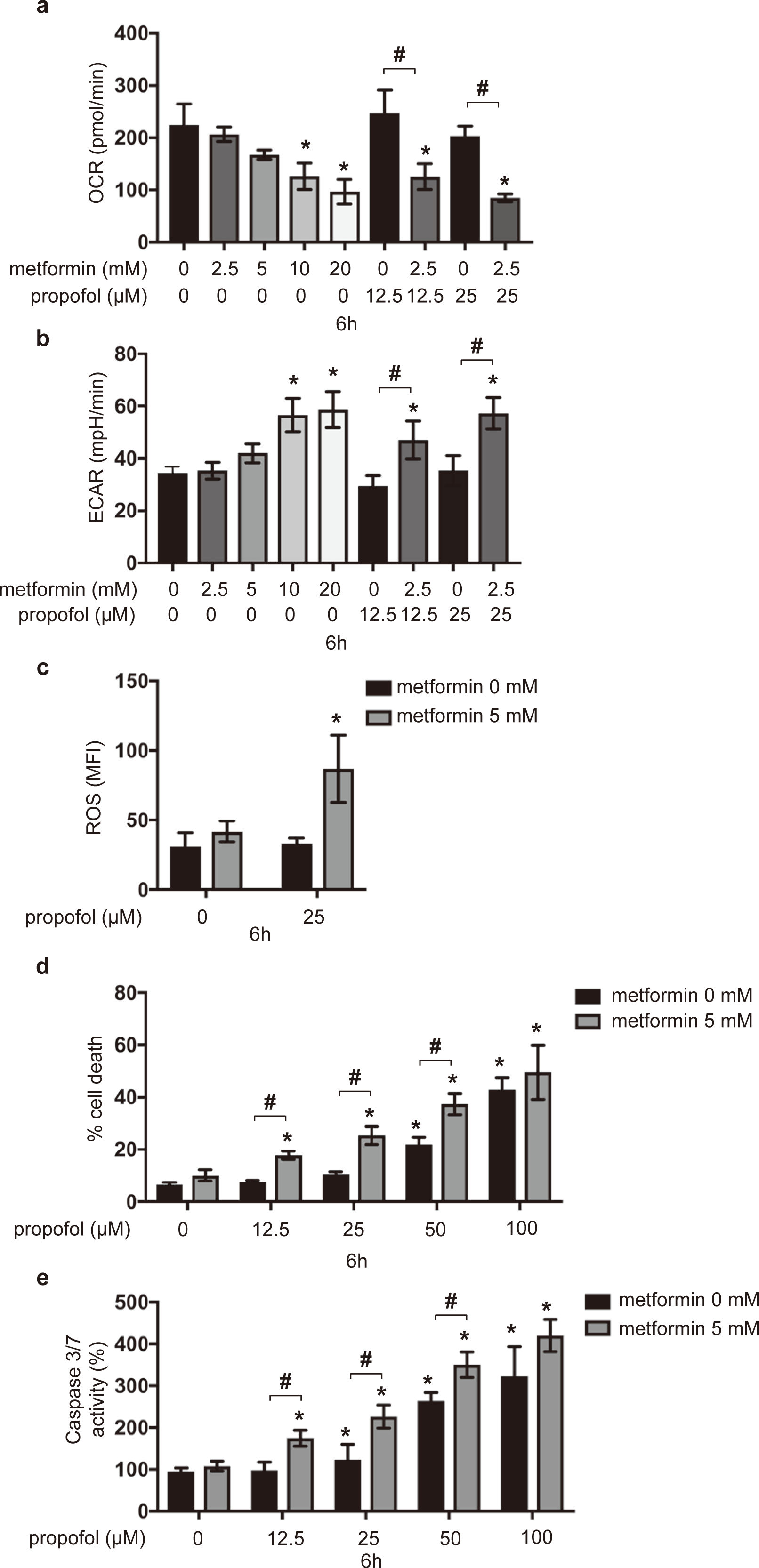
Synergistic effects of propofol and metformin on caspase activity and cell death. (a) OCR and (b) ECAR of SH-SY5Y cells exposed to the indicated concentrations of metformin (2.5, 5, 10, and 20 mM) for 6 h. (c) SH-SY5Y cells were exposed to 25 μM propofol with or without 5 mM metformin, and ROS production was determined (n = 3). (d and e) SH-SY5Y cells were exposed to the indicated concentrations (12.5, 25, 50, and 100 μM) of propofol with or without 5 mM metformin for 6 h. (d) Cells were harvested, and percentages of cell death were measured by flow cytometry. The ratio of PI-positive and/or annexin V-positive cells [(Q1 + Q2 + Q4)/(Q1 + Q2 + Q3 + Q4)] was used to calculate the percentage of dead cells (n = 3). (e) Caspase-3/7 activity in each treatment group (n = 3). Data presented in (a–e) are expressed as the means ± SD. Differences between treatment groups were evaluated by one-way ANOVA, followed by Dunnett’s multiple comparison test (a, b, and c), or by two-way ANOVA, followed by Dunnett’s multiple comparison test (d and e). \**p* < 0.05 compared to the control cell population; #*p* < 0.05 compared to the indicated experimental groups.

## Discussion

In this study, we demonstrated for the first time that propofol at clinically relevant concentrations and within clinically relevant incubation times altered the oxygen metabolism by targeting mitochondrial complexes I and III and induced a cellular metabolic switch, from OXPHOS to glycolysis, and generation of ROS. The suppression of mitochondria elicited the cell death in cell lines of various origins, including transmitochondrial cybrids carrying mtDNA with defined pathogenic mutations.

The concentrations of propofol tested in this study varied from 12.5 to 150 μM. It has been reported that the plasma concentrations of propofol during anesthesia and sedation are from 2 μg/mL (11 μM) up to 5 μg/mL (27.5 μM) ^23^. The concentration of propofol was also measured in tissue samples from rats treated with propofol at a dose of 20 mg/kg/h, and the study indicated that the tissue concentration of propofol could reach 200 μM under certain conditions ^24^. Thus, the evidence indicates that the concentrations of propofol used in this study were clinically relevant. The duration of exposure used in this study ranged from 3 to 12 h, which was also within the clinically relevant period of exposure.

Although predictive factors for PRIS have not been established, there is a consensus that exposure to high doses of propofol for prolonged periods is the most critical risk factor for PRIS ^1,2,6,25^. In this study, we demonstrated that propofol at concentrations equal to or greater than 50 μM but not at or below 25 μM showed cell toxicity within 6 h. On the other hand, even 25 μM propofol significantly increased the cell death and caspase activity during incubation for 12 h (Figs. 1b and 1f). In addition, propofol at 50 and 25 μM significantly suppressed OCR and increased ECAR within 6 and 12 h, respectively. At 100 μM, propofol inhibited both OCR and ECAR, which could be due to a rapid cell death induced by 100 μM propofol. The evidence indicates that propofol at clinically relevant concentrations suppresses OXPHOS and induces a metabolic switch, from OXPHOS to glycolysis, resulting in the enhancement of lactate production. This metabolic conversion can be one of the most critical cellular mechanisms of lactic acidosis observed in PRIS ^1,2^.

As clearly demonstrated in this and other studies ^26-29^, propofol induces cell death. In this study, we demonstrated that not only neuronal SH-SY5Y cells but also cells of other origins, such as C2C12 muscle cells, HeLa cervical carcinoma cells, and P29 lung cancer cells, are susceptible to propofol. Although there is no consensus on the target organs or tissues in the context of PRIS, our results suggest that propofol can exert toxicity against a wide range of tissues.

There are at least two known modes of cell death, apoptosis and necrosis. Apoptosis is a strictly regulated or programmed process involving the activation of specific proteases, which are responsible for organized removal of damaged cells ^30^. Necrosis is a pathophysiologically different form of cell death, which is accompanied by a loss of Δψm and impairment of OXPHOS. In this study, propofol treatment promoted the caspase-9 and caspase-3/7 activation. These data indicate that propofol at least activates the apoptosis pathway. Meanwhile, the flow cytometry analysis demonstrated that treatment of cells with propofol concentrations greater than 50 μM for 12 h also resulted in an LDH release (Fig. 1d) and increased the PI^+^/annexin V^−^ cell population, in addition to the PI^+^/annexin V^+^ cell population (Fig. 1a). These data strongly suggest that propofol elicits both types of cell death, resulting in cell membrane injury at a 50 μM concentration and 12-h exposure.

As demonstrated in figures 2 and 3, propofol inhibited the mitochondrial oxygen consumption in a time‐ and concentration-dependent manner. In this study, we measured OCR, which is dependent on the activity of each ETC complex in intact cells, using an extracellular flux analyzer. Several studies have used isolated mitochondria, and some have suggested that propofol inhibits the enzymatic activity of complexes II and IV ^31^. However, our study using intact cells indicated that propofol does not affect the complex II‐ or complex IV-dependent OCR. Another finding was that propofol induced ROS generation (Fig. 2g). While the precise origin of ROS remains unclear, our findings strongly suggest that mitochondria play a critical role in this process. ρ0P29 cells and P29mtΔ cells did not produce ROS in response to propofol treatment (data not shown). It has been demonstrated that the disturbance of mitochondria leads to increased production of ROS by ETC ^12,13,32-34^. In fact, cybrid cells such as P29mtA11 and P29mtB82M with a mutation in the mitochondrial NADH dehydrogenase subunit 6 (*ND6*) gene generate ROS under 20% O_2_ conditions ^16,35^, which implies that propofol-mediated cell death is dependent on ROS from mitochondria. Mitochondrial disease, once thought to be a rare clinical entity, is now recognized as an important cause of a wide range of neurological, cardiac, muscle, and endocrine disorders ^4,5^. It has been reported that complex I is uniquely sensitive to many anesthetic agents ^36^. To approach the issue, we used cells harboring mtDNA mutations ^15,16,37^. As shown in figures 3 and 6, ρ0P29 cells, which do not have mtDNA, are resistant to 50 μM propofol. P29mtΔ cells, carrying the nuclear genome of P29 cells and the mitochondrial genome of ΔmtDNA4696, with a 4,696-bp deletion ^15^, are also resistant to propofol. In contrast, P29mtA11 cells with the G13997A mutation in mtDNA and P29mtB82M cells with the 13885insC insertion, both of which affect the NADH-ubiquinone oxidoreductase chain 6 (ND6) protein of complex I, are more sensitive to propofol than P29 cells are. Propofol at 12.5 and 25 μM induced the cell death and caspase-3/7 activation in both mutant cell lines within 6 h (Fig. 6c). This is consistent with the evidence showing that 25 μM propofol induced the cell death in the presence of a sublethal concentration of rotenone. P29mtCOIM cells with a missense mutation in cytochrome c oxidase I (COI) are as sensitive to propofol as P29mtA11 and P29mtB82M cells. This is also consistent with the result showing that propofol induced the cell death in the presence of oligomycin (Fig. 5). This evidence indicates that cells with mutations in mtDNA are more sensitive to propofol.

In addition to the genetic mutations affecting mitochondrial function, pharmacological disturbance of mitochondria with biguanides, including metformin and phenformin, also synergistically suppressed the mitochondrial function and induced caspase activation and cell death. The primary effect of biguanides is generally thought to be the inhibition of respiratory complex I (NADH:ubiquinone oxidoreductase), which leads to the energy stress by decreasing ATP synthesis by OXPHOS ^18,19,38,39^. In addition, a number of studies have indicated that biguanides also affect complexes II, III, and IV and F1F0-ATPase ^20,40^. Metformin and phenformin are used as antidiabetic drugs and are associated with lactic acidosis ^20,41,42^. As demonstrated in this study, both metformin and phenformin increased ECAR and decreased OCR, which indicates a metabolic switch. In addition to biguanides, other clinically used drugs, including chloramphenicol ^43,44^, aspirin ^45,46^, statins ^18^, and local anesthetics ^47^, also inhibit mitochondrial functions. Our results indicate that pre-exposure to mitochondrial inhibitors may increase the toxicity of propofol.

There are some limitations in this study. Thus, we only used cultured cells. Because PRIS is a systemic syndrome, using cells is not sufficient to completely understand the pathophysiology of PRIS. An *in vivo* investigation using experimental animals may be warranted to confirm our findings.

In conclusion, clinically relevant concentrations of propofol used within a clinically relevant exposure time suppressed the mitochondrial function, caused the generation of ROS, and induced metabolic reprogramming, from OXPHOS to glycolysis, by targeting complexes I and III of mitochondria. Our data also indicated that a predisposition to mitochondrial dysfunction, caused by genetic mutations and pharmacological suppression of ETC by biguanides such as metformin and phenformin, promoted the cell death and caspase activation by propofol. The process is likely to constitute the molecular basis of PRIS.

## Materials and Methods Reagents

Propofol (2,6-diisopropylphenol), 2,4-diisopropylphenol, and dimethyl sulfoxide were obtained from Sigma–Aldrich (St. Louis, MO, USA). Rotenone, oligomycin, and antimycin A were obtained from Abcam, Inc. (Cambridge, MA, USA). The Key Resources Table is provided as Table 1.

**Table 1.**
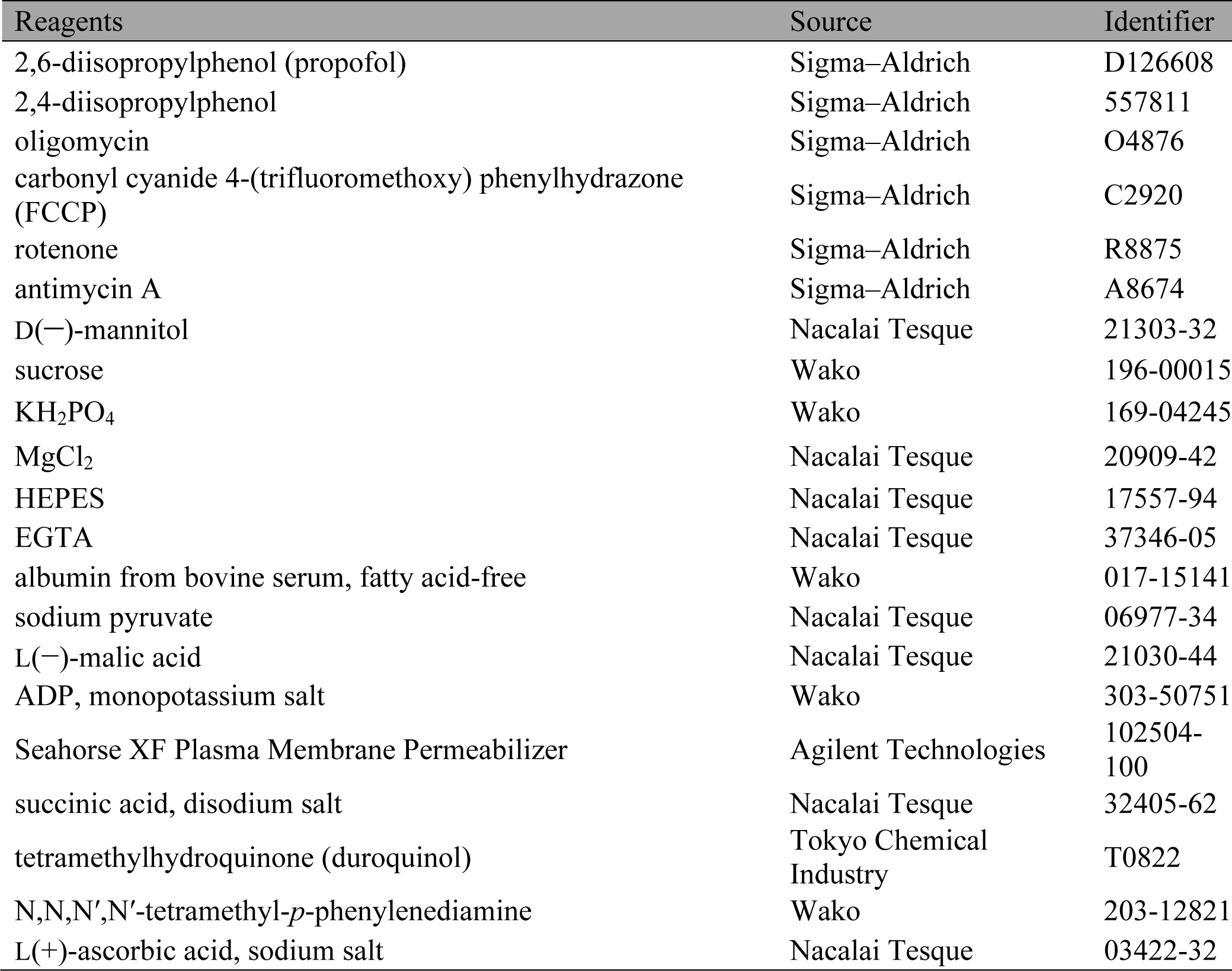
Key Resources Table.

### Cell lines and cell culture

Established cell lines derived from human neuroblastoma SH-SY5Y cells and cervical carcinoma HeLa cells were maintained in RPMI 1640 medium supplemented with 10% fetal bovine serum, 100 U/mL penicillin, and 0.1 mg/mL streptomycin. The mouse cell lines and their characteristics are listed in Tables 2 and 3. P29 cells originated from Lewis lung carcinoma (C57BL/6 mouse strain), and B82 cells are fibrosarcoma cells derived from the L929 fibroblast cell line (C3H/An mouse strain) ^15,16,48^. Parental P29 cells, ρ0 cells, and the transmitochondrial cybrids were grown in Dulbecco’s modified Eagle’s medium (DMEM) supplemented with pyruvate (0.1 mg/mL), uridine (50 mg/mL), and 10% fetal bovine serum. We isolated ρ0 cells by treating parental P29 cells with 1.5 mg/mL ditercalinium, an antitumor *bis*-intercalating agent. Enucleated cells of mtDNA donors were prepared by pretreatment with cytochalasin B (10 μg/mL) for 2 min, followed by centrifugation at 7,500 × *g* for 10 min ^16^. The resultant cytoplasts were fused with ρ0 cells using polyethylene glycol. The transmitochondrial cybrids (see Table 3) were isolated in a selection medium that allows exclusive growth of the cybrids^16^.

**Table 2.**
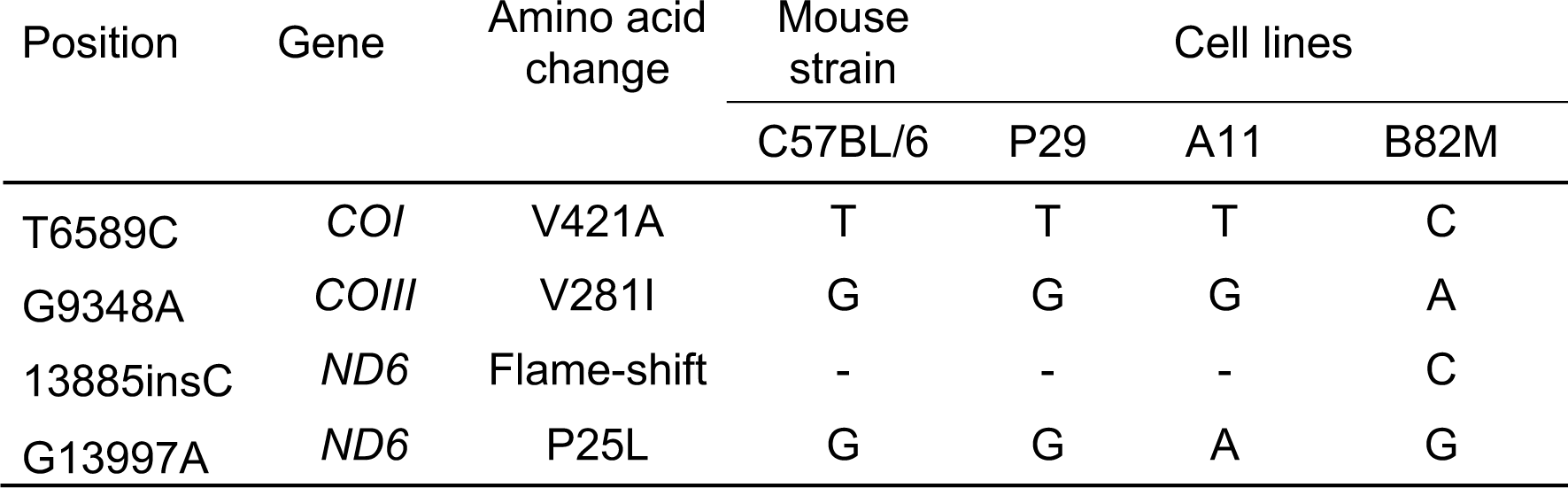
Identification of pathogenic mutations in mtDNA sequences.

The G13997A mutation in *ND6* is a missense mutation that changes the amino acid proline to leucine at a site that is highly conserved throughout vertebrates. The 13885insC mutation in *ND6* is a frameshift mutation that has been previously reported as a pathogenic mutation inducing substantial complex I defects in some sublines of the L929 fibroblast cell line.

Bases: A, Adenine; G, Guanine; C, Cytosine; T: Thymine

Amino acids: V, Valine; A, Alanine; I, Isoleucine; P, Proline; L, Leucine

**Table 3.** Genetic characteristics of parent cells and their transmitochondrial cybrids.

| Cell lines | Nuclear<br>genotype | Fusion combinations |  |
| --- | --- | --- | --- |
|  |  | Nuclear<br>donor | mtDNA donors |
| ρ0P29 | P29 | ρ0P29 | (–) |
| P29mtA11 | P29 | ρ0P29 | enA11 |
| P29mtB82M | P29 | ρ0P29 | enB82M |
| P29mtCOIM | P29 | ρ0P29 | enB82 |
| P29mtΔ | P29 | ρ0P29 | enB82mtΔ |

ρ0, mtDNA-less cells; en, enucleated donor.

### Cell growth MTS assay

Cell growth was assessed using a CellTiter 96 AQueous One Solution Cell Proliferation Assay™ with MTS (Promega, Madison, WI, USA) ^47,49^. Briefly, cells were seeded into 96-well plates (2 × 10^4^cells/well) and cultured overnight. On the following day, the cells were treated with the indicated concentrations of the appropriate drug(s) for varying times. After the treatment, 20 μL of the CellTiter 96 AQueous One Solution™ reagent was added to each well, the plates were incubated at 37 °C for 1 h, and the absorbance of each sample was measured using an iMark™ microplate reader (Bio-Rad, Hercules, CA, USA) at a wavelength of 490 nm. Cell viability was calculated by comparing the absorbance of treated cells with that of the control cells incubated without drugs, which was defined as 100%. All samples were tested in triplicate or quadruplicate in each experiment.

### Caspase-3/7 and caspase-9 activity assays

Activity levels of caspase-3/7 and caspase-9 were assessed using an Apo-ONE™ homogeneous caspase-3/7 assay kit (Promega) and a Caspase-Glo™ 9 assay kit (Promega), respectively, according to the manufacturer’s protocols ^47,49^. Briefly, cells were seeded into 96-well plates (2 × 10^4^cells/well) and incubated overnight. On the following day, the cells were treated with the indicated concentrations of the appropriate drug(s) for varying times. After the treatment, 100 μL of the Apo-ONE™ caspase-3/7 reagent was added to each well. The plates were incubated at room temperature for 1 h, and the luminescence of each well was measured using an EnSpire™ multimode plate reader (PerkinElmer, Waltham, MA, USA). Caspase activity was calculated by comparing the levels of luminescence of the treated cells with that of the control cell population incubated without drugs, which was defined as 100%. The assays were performed in triplicate at least twice. Data were expressed as the mean ± standard deviation (SD).

### Analysis of cell death

The protocol was described previously ^47,49^. Briefly, the levels of cell apoptosis were measured using an Annexin V–FITC apoptosis detection kit (BioVision, Milpitas, CA, USA), according to the manufacturer’s instructions. For the analysis, cells were seeded into 6-well plates (3 × 10^5^cells/well) and incubated overnight. On the following day, the cells were treated with the indicated concentrations of the appropriate drug(s) for varying times and harvested by centrifugation at 1,200 rpm for 3 min. The cell pellets were resuspended in a mixture comprised of 500 μL of binding buffer, 5 μL of annexin V–FITC, and 5 μL of PI (50 μg/mL). The suspensions were incubated for 5 min at room temperature in the dark and analyzed using a FACSCalibur flow cytometer (BD Biosciences, San Jose, CA, USA) equipped with the CellQuest Pro™ software. The data were evaluated using the FlowJo™ version 9.9.4 software (TreeStar, Ashland, OR, USA), then exported to Excel spreadsheets, and subsequently analyzed using the statistical application GraphPad™ Prism 7.

### Determination of mitochondrial membrane potential

The mitochondrial membrane potential (ΔΨm) was determined by flow cytometry using a MitoPT™ JC-1 assay kit (ImmunoChemistry Technologies, Bloomington, MN, USA), according to the manufacturer’s instructions ^49^. For the analysis, cells were seeded into 6-well plates (3 × 10^5^cells/well) and cultured overnight. On the following day, the cells were treated with the indicated concentrations of the appropriate drug(s) for varying times and then pelleted by centrifugation at 1,200 rpm for 3 min. The cells were then resuspended in JC-1, incubated at 37 °C for 15 min in the dark, and collected by centrifugation at 1,200 rpm for 3 min. The cell pellets were resuspended in 500 μL of assay buffer. The samples were subsequently analyzed using a FACSCalibur flow cytometer (BD Biosciences) equipped with the CellQuest Pro™ software for the detection of red JC-1 aggregates (590 nm emission) or green JC-1 monomers (527 nm emission). The data were evaluated using the FlowJo version 7.6.3 software (TreeStar), then exported to Excel spreadsheets, and subsequently analyzed using the statistical application GraphPad™ Prism 7.

### LDH-based cytotoxicity assay

Cytotoxicity was measured using a CytoTox-ONE™ kit (Promega) as described previously ^47,49^. Briefly, cells were cultured overnight in 96-well plates (2 × 10^4^cells/well) and treated with the indicated drug(s) for varying lengths of time. Twenty microliters of the CytoTox-ONE™ reagent was added to each well, and the plates were incubated at 22 °C for 10 min. The reaction was terminated by adding 50 μL of the stop solution, and the fluorescence was recorded at an excitation wavelength of 560 nm and an emission wavelength of 590 nm using an EnSpire™ multimode plate reader (PerkinElmer). The percentage of cell death was determined by comparing the release of LDH (fluorescence value) in each treatment group with that of the positive control treated with the lysis solution, which was defined as 100%. Meanwhile, the level of LDH release from untreated cells (negative control) was defined as 0%. Each sample was assayed in triplicate.

### Measurement of ROS generation

ROS generation was detected with 2′,7′-dichlorofluorescin diacetate (DCFH-DA) (Molecular Probes, Eugene, OR, USA). Briefly, cells cultured in 35-mm-diameter glass-bottom culture dishes (MatTek, Ashland, MA, USA) were incubated with 10 μM DCFH-DA for 10 min at 37 °C in a serum-free DMEM, then washed twice with Dulbecco’s phosphate-buffered saline, and analyzed using a flow cytometer (Beckton Dickinson). The mean fluorescence intensity was analyzed using the CellQuest software (Becton Dickinson).

### Cellular oxygen consumption and extracellular acidification measurement

Cellular OCR and ECAR were determined with the XF Cell Mito Stress Test™ and XF Glycolysis Stress Test™, respectively, using an XFp Extracellular Flux Analyzer™ (Seahorse Bioscience, USA) ^47^. Cells (2 × 10^5^cells/well) were seeded into an XFp cell culture microplate, and OCR was assessed in glucose-containing XF base medium according to the manufacturer’s instructions. The sensor cartridge for the XFp analyzer was hydrated in a 37 °C non-CO_2_ incubator on the day before the experiment. For the OCR assay, injection port A on the sensor cartridge was loaded with 1.5 μM oligomycin (complex V inhibitor), port B was loaded with 2 μM carbonyl cyanide-4-(trifluoromethoxy) phenylhydrazone (FCCP), and port C was loaded with 0.5 μM rotenone/antimycin A (inhibitors of complex I and complex III). During the sensor calibration, cells were incubated in a 37 °C non-CO_2_ incubator in 180 μL of assay medium (XF base medium with 5.5 mM glucose, 1 mM pyruvate, and 2 mM L-glutamine, pH 7.4). The plate was immediately placed into the calibrated XFp extracellular flux analyzer for the Mito Stress test (Supplementary Fig. 3a). The assay parameters were calculated as follows: OCR (basal) = (last rate measurement before oligomycin injection) − (minimum rate measurement after rotenone/antimycin-A injection); OCR (maximal) = (maximum rate measurement after FCCP injection) − (minimum rate measurement after rotenone/antimycin A injection); OCR (non-mitochondrial respiration) = (minimum rate measurement after rotenone/antimycin A injection); proton leak = (minimum rate measurement after oligomycin injection) – (non-mitochondrial respiration). For the ECAR assay, injection port A on the sensor cartridge was loaded with 10 mM glucose. During the sensor calibration, cells were incubated in a 37 °C non-CO_2_ incubator in 180 μL of assay medium (XF base medium with 2 mM L-glutamine, pH 7.4). The plate was immediately placed into the calibrated XFp Extracellular Flux Analyzer™ for the Glycolysis Stress test (Supplementary Fig. 3b). Oligomycin (1 μM) and 50 mM 2-deoxy-D-glucose were loaded for the measurement. ECAR was normalized for the total protein/well and calculated as follows: ECAR (glycolysis) = (maximum rate measurement after glucose injection) − (last rate measurement before glucose injection).

### Measurement of oxygen consumption in permeabilized cells

The activity of individual respiratory chain complexes was evaluated in permeabilized cells ^50,51^. Briefly, cells were washed with MAS buffer (220 mM mannitol, 70 mM sucrose, 10 mM KH_2_PO_4_, 5 mM MgCl_2_, 2 mM HEPES, 1 mM EGTA, 0.2% fatty acid-free bovine albumin, adjusted to pH 7.2 with KOH), and the medium was replaced with MAS buffer supplemented with 10 mM pyruvate, 1 mM malate, 4 mM ADP, and 1 nM plasma membrane permeabilizer^™^. The cells were then loaded into the XFp analyzer to measure respiration rates using cycles of 30 s mixing/30 s waiting/2 min measurement. After measurement of pyruvate-driven respiration, rotenone (final concentration: 2 μM) was injected through port A to halt the complex I-mediated respiratory activity. Next, succinate (10 mM) was injected through port B to donate electrons at complex II, bypassing complex I inhibition. Addition of antimycin A (2 μM) via port C inhibited complex III, and N,N,N′,N′-tetramethyl-*p*-phenylenediamine (0.1 mM), combined with ascorbate (10 mM), was subsequently injected through port D to measure complex IV activity. As an alternative approach, cells were initially provided with pyruvate to measure complex I activity. After injection of rotenone, duroquinol was injected to stimulate complex III-mediated respiration.

### Statistical analysis

All experiments were repeated at least twice, and each sample was evaluated in triplicate. Representative data, expressed as the mean ± SD, are shown. Differences between treatment groups were evaluated by one-way analysis of variance (ANOVA) or two-way ANOVA, followed by Dunnett’s multiple comparison test or *t*-test using GraphPad Prism 7. *P*-values of < 0.05 were considered statistically significant.

## Acknowledgments

This work was supported by the Japan Society for the Promotion of Science KAKENHI, grants #26670693 and #24592336 to K.H., #25462457 to K.N., and #15K15577 to T.A, a research grant from Katano Kai to K.H. and A.O., research grant B from Kansai Medical University to A.O., and a KMU consortium grant from Kansai Medical University to K.H.

We would like to thank Editage (<u>www.editage.jp</u>) for English language editing.

## Author contributions statement

C.S. performed the experiments, analyzed the data, and co-wrote the manuscript. A.O., M.T., M.K., H.S., T.U., Y.M., and T.A. contributed to the data analysis and discussion. J.H. and K.T. provided experimental materials and contributed to the discussion. K.H. designed and supervised the study, analyzed the data, and co-wrote the manuscript.

## Additional information Competing financial interests

The authors declare no competing financial interests.

## Supplementary Information

**Supplementary Figure 1.**
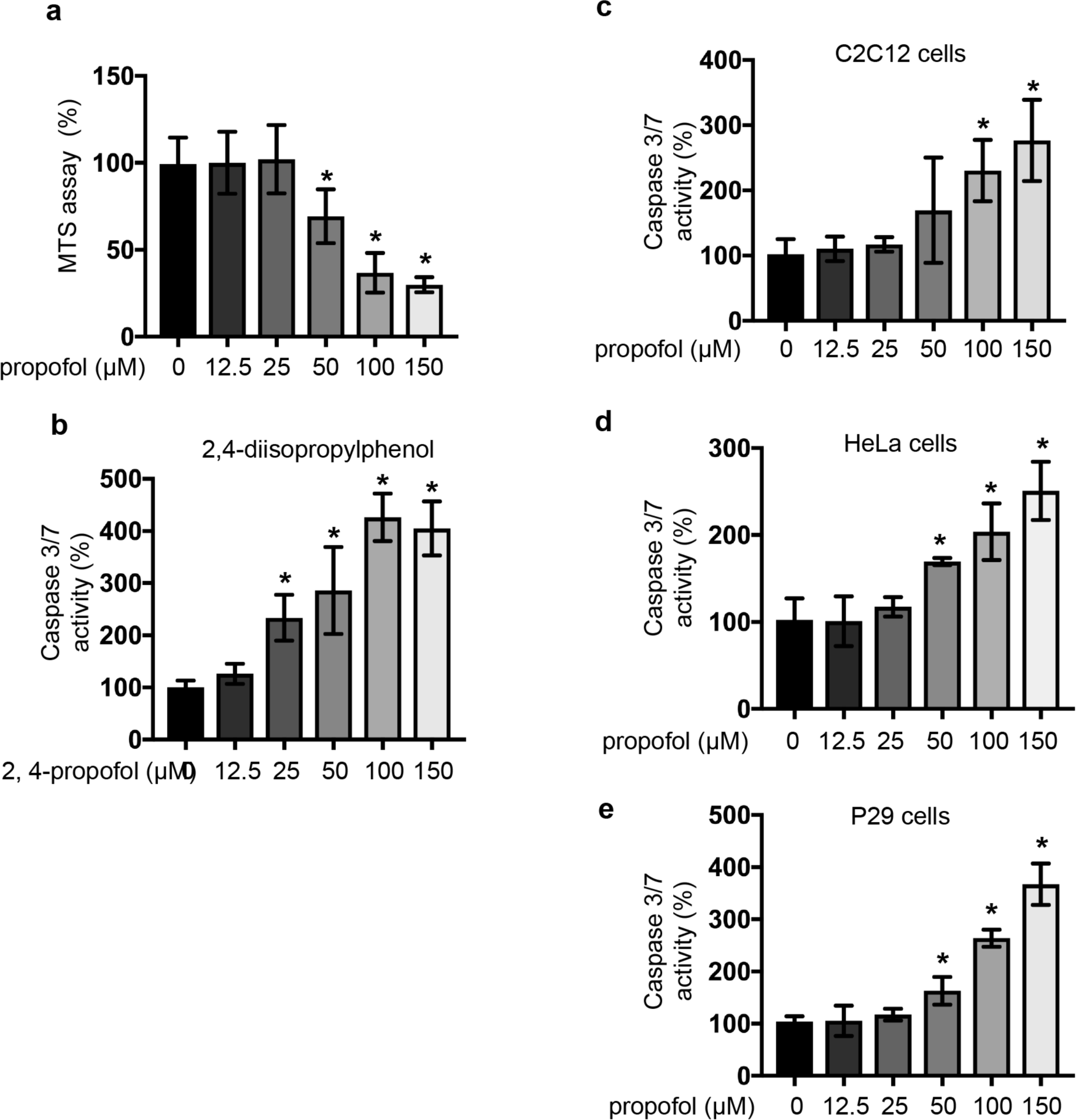
Analysis of cell proliferation, caspase 3/7 activity. (a) SH-SY5Y cells were exposed to the indicated concentrations (12.5, 25, 50, 100 or 150 μM) of propofol for 6 h. Graphic depiction of the levels of cell proliferation of treated and untreated cells, as evaluated by MTS [3-(4,5-dimethylthiazol-2-yl)-5-(3-carboxymethoxyphenyl)-2-(4-sulfophenyl)-2H-tetrazolium] assay (n = 3) as described in Cell growth MTS assay section in Materials and Methods section. (b) SH-SY5Y cells were exposed to the indicated concentrations (12.5, 25, 50, 100 and 150 μM) of 2,4 diisopropylphenol 6 h. Graphic depictions of caspase-3/7 (n = 3). (c, d and e) C2C12 cells (c), HeLa cells (d) and P29 cells (e) were exposed to the indicated concentrations (12.5, 25, 50, 100 and 150 μM) of propofol for 6 h. Graphic depictions of caspase-3/7 activity (n = 3). Differences between treatment groups were evaluated by one-way ANOVA, followed by Dunnett’s multiple comparison test. *p < 0.05 compared to the control cell population (incubation for 0 h, no treatment).

**Supplementary Figure 2.**
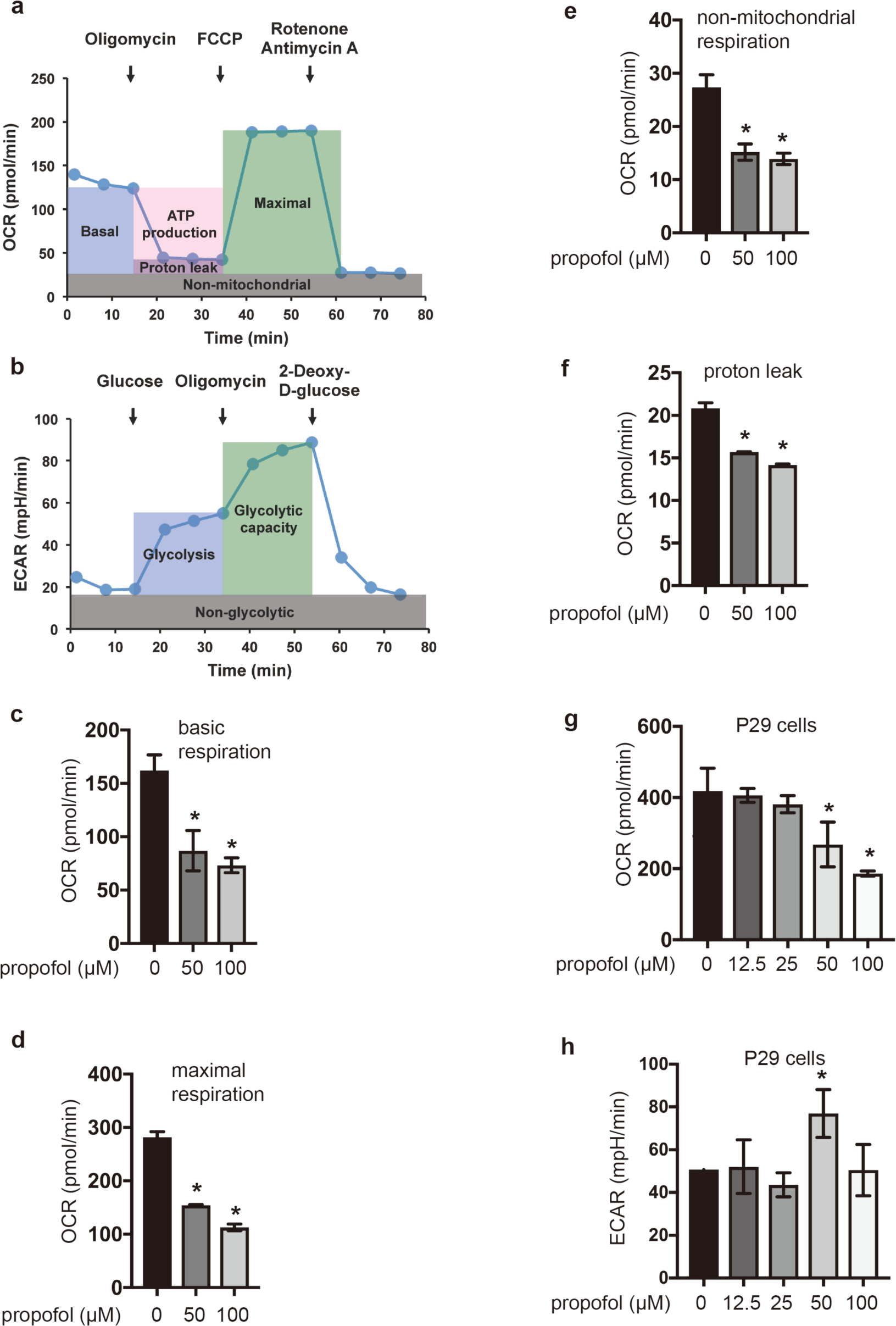
Oxygen metabolism and ROS generation in SH-SY5Y cells treated with propofol. (a) Cell Mito Stress test profile of the key parameters of mitochondrial oxygen consumption rate (OCR). (b) Cell glycolysis test profile of the key parameters of the extracellular acidification rate (ECAR). (c, d, e and f) Sequential compound injections measure basal respiration, maximal respiration, and non-mitochondrial respiration, non-mitochondrial repiration and proton leak were demonsrated. OCR (basal respiration) (c) OCR (maximal respiration) (d), OCR (non-mitochondrial respiration) (e) and Proton Leak (f) in SH-SY5Y cells treated with indicated doses of propofol were demonstrated. (g and h) OCR (f) and ECAR (h) in P29 cells exposed to the indicated doses of propofol (12.5, 25, 50 and 100 μM) for 6h. Data presented are expressed as means ± standard deviations (SD). Differences between results were evaluated by the one-way ANOVA followed by Dunnett’s multiple comparison test *p < 0.05 compared to the control cell population.

**Supplementary Figure 3.**
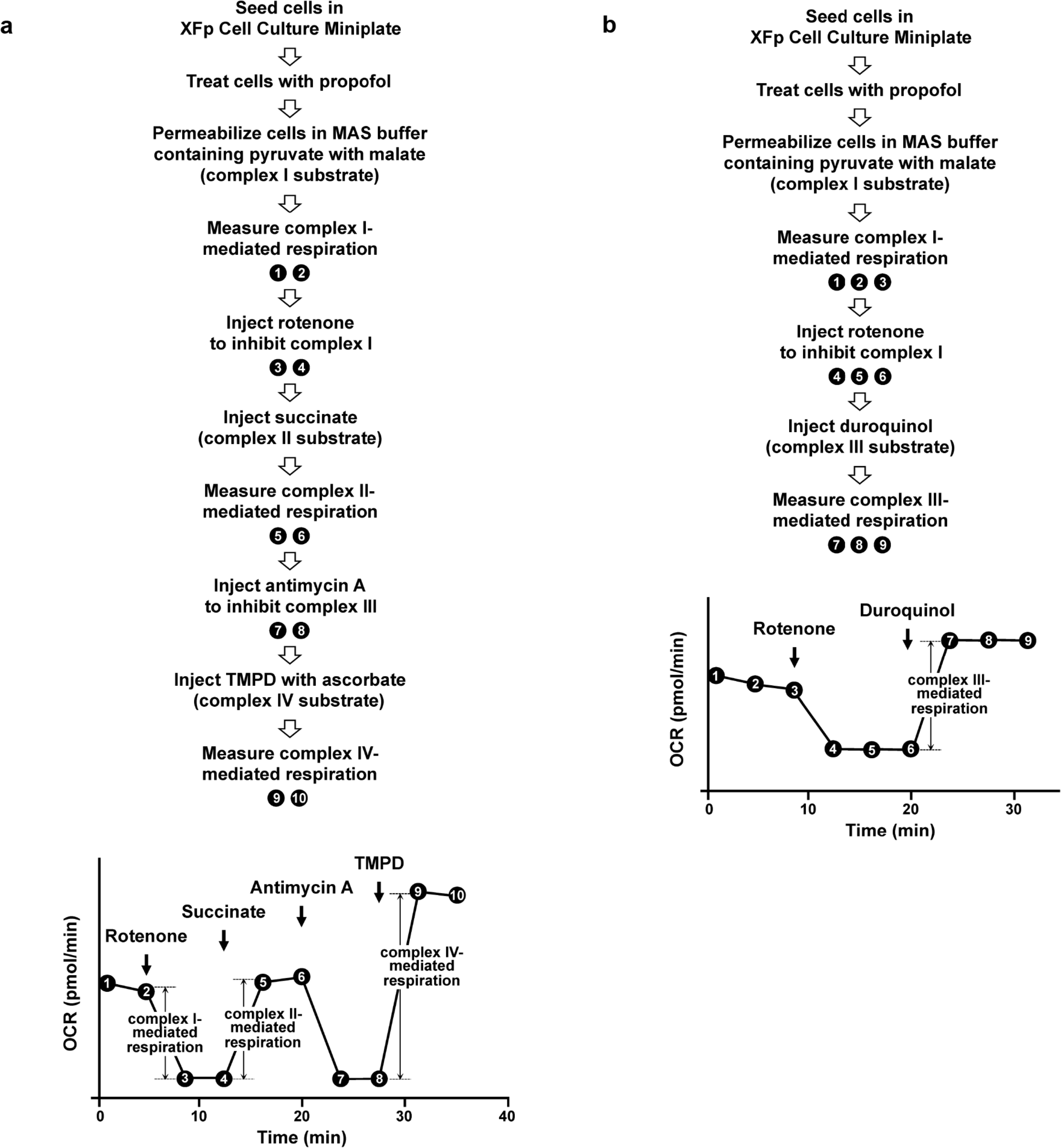
Measurement of oxygen consumption in permeabilized cells. The activity of individual respiratory chain complexes was evaluated by employing specific substrates and inhibitors. (a) Cells were treated with plasma membrane permeabilizer and offered pyruvate and malate for measuring complex I-mediated respiration. Cells were sequentially treated with rotenone (complex I inhibitor), succinate (complex II substrate), antimycin A (complex III inhibitor), and TMPD plus ascorbate (complex IV substrate) as indicated. Oxygen consumption measurements were performed using a Seahorse XFp Extracellular Flux Analyzer. Distinct complex activities were calculated as follows: complex I-mediated respiration = (mean OCR value between points 1 and 2) - (mean OCR value between points 3 and 4); complex II-mediated respiration = (mean OCR value between points 5 and 6) - (mean OCR value between points 3 and 4); complex IV-mediated respiration = (mean OCR value between points 9 and 10)-(mean OCR value between points 7 and 8). (b) Cells were permeabilized as in (a), and treated with rotenone, followed by duroquinol to donate electrons at complex III. Complex III-mediated respiratory activity was calculated as (mean OCR value between points 7-9) - (mean OCR value between points 4-6).

**Supplementary Figure 4.**
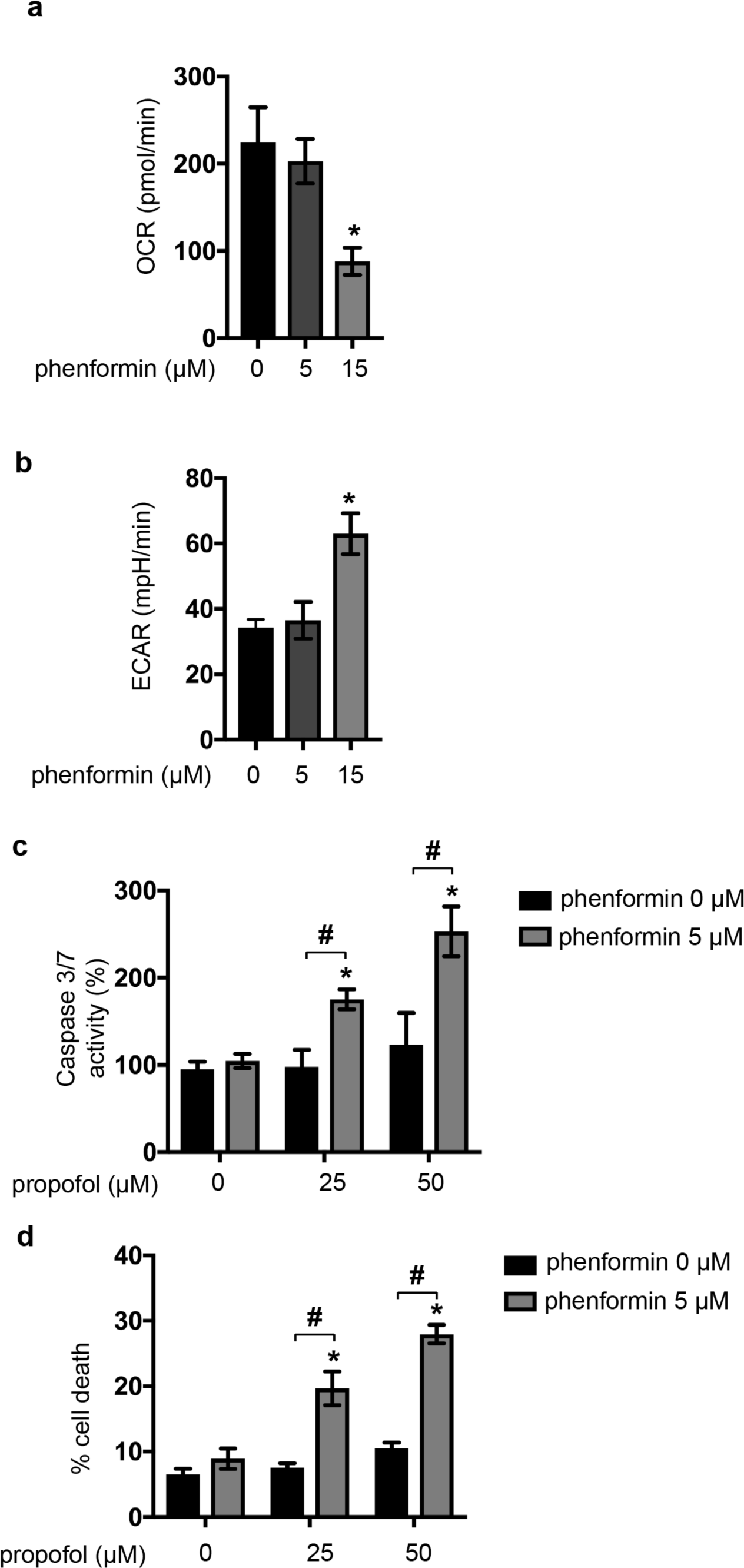
Synergistic effect of propofol with the biguanide phenformin on caspase activity and cell death. Oxygen consumption rate (OCR) (a) and extracellular acidification rate (ECAR) (b) of SH-SY5Y cells exposed to the indicated doses of phenformin (5 and 15 μM) for 6h. SH-SY5Y cells were exposed to the indicated concentrations (25 and 50 μM) of propofol with or without treatment with 5 μM phenformin for 6 h. (c) Graphic depictions of caspase-3/7 (n = 3) activity in each treatment group. (n=3) (d) Cells were harvested and cell death percentages were measured with flow cytometry. The ratio of propidium iodide (PI)-positive and/or annexin V-positive cells [(Q1 + Q2 + Q4)/(Q1 + Q2 + Q3 + Q4)] was used to calculate the percentage of dead cells (n = 3). Data presented in (a)-(d) are expressed as means ± standard deviations (SD). Differences between results were evaluated by the one-way ANOVA followed by Dunnett’s multiple comparisons test (a and b), and two-way ANOVA followed by Dunnett’s multiple comparisons test (c and d). *p < 0.05 compared to the control cell population.

## References

1 Kam, P. C. & Cardone, D. Propofol infusion syndrome. Anaesthesia 62, 690-701, doi:10.1111/j.1365-2044.2007.05055.x (2007).

2 Bray, R. J. Propofol infusion syndrome in children. Paediatr Anaesth 8, 491-499 (1998).

3 Parke, T. J. et al. Metabolic acidosis and fatal myocardial failure after propofol infusion in children: five case reports. BMJ 305, 613-616 (1992).

4 Finsterer, J. & Frank, M. Propofol Is Mitochondrion-Toxic and May Unmask a Mitochondrial Disorder. J Child Neurol 31, 1489-1494, doi:10.1177/0883073816661458 (2016).

5 Niezgoda, J. & Morgan, P. G. Anesthetic considerations in patients with mitochondrial defects. Paediatr Anaesth 23, 785-793, doi:10.1111/pan.12158 (2013).

6 Fudickar, A. & Bein, B. Propofol infusion syndrome: update of clinical manifestation and pathophysiology. Minerva Anestesiol 75, 339-344 (2009).

7 Bains, R., Moe, M. C., Vinje, M. L. & Berg-Johnsen, J. Sevoflurane and propofol depolarize mitochondria in rat and human cerebrocortical synaptosomes by different mechanisms. Acta Anaesthesiol Scand 53, 1354-1360, doi:10.1111/j.1399-6576.2009.02047.x (2009).

8 Branca, D., Roberti, M. S., Vincenti, E. & Scutari, G. Uncoupling effect of the general anesthetic 2,6-diisopropylphenol in isolated rat liver mitochondria. Arch Biochem Biophys 290, 517-521 (1991).

9 Rigoulet, M., Devin, A., Averet, N., Vandais, B. & Guerin, B. Mechanisms of inhibition and uncoupling of respiration in isolated rat liver mitochondria by the general anesthetic 2,6-diisopropylphenol. Eur J Biochem 241, 280-285 (1996).

10 Cray, S. H., Robinson, B. H. & Cox, P. N. Lactic acidemia and bradyarrhythmia in a child sedated with propofol. Crit Care Med 26, 2087-2092 (1998).

11 Prabhakar, N. R. & Semenza, G. L. Adaptive and maladaptive cardiorespiratory responses to continuous and intermittent hypoxia mediated by hypoxia-inducible factors 1 and 2. Physiol Rev 92, 967-1003, doi:10.1152/physrev.00030.2011 (2012).

12 Murphy, M. P. How mitochondria produce reactive oxygen species. Biochem J 417, 1-13, doi:10.1042/BJ20081386 (2009).

13 Chen, Q., Vazquez, E. J., Moghaddas, S., Hoppel, C. L. & Lesnefsky, E. J. Production of reactive oxygen species by mitochondria: central role of complex III. J Biol Chem 278, 36027-36031, doi:10.1074/jbc.M304854200 (2003).

14 Chandel, N. S. & Schumacker, P. T. Cells depleted of mitochondrial DNA (rho0) yield insight into physiological mechanisms. FEBS Lett 454, 173-176 (1999).

15 Ishikawa, K. et al. Enhanced glycolysis induced by mtDNA mutations does not regulate metastasis. FEBS Lett 582, 3525-3530, doi:10.1016/j.febslet.2008.09.024 (2008).

16 Ishikawa, K. et al. ROS-generating mitochondrial DNA mutations can regulate tumor cell metastasis. Science 320, 661-664, doi:10.1126/science.1156906 (2008).

17 Ferrannini, E. The target of metformin in type 2 diabetes. N Engl J Med 371, 1547-1548, doi:10.1056/NEJMcibr1409796 (2014).

18 Wheaton, W. W. et al. Metformin inhibits mitochondrial complex I of cancer cells to reduce tumorigenesis. Elife 3, e02242, doi:10.7554/eLife.02242 (2014).

19 Gui, D. Y. et al. Environment Dictates Dependence on Mitochondrial Complex I for NAD+ and Aspartate Production and Determines Cancer Cell Sensitivity to Metformin. Cell Metab 24, 716-727, doi:10.1016/j.cmet.2016.09.006 (2016).

20 Bridges, H. R., Jones, A. J., Pollak, M. N. & Hirst, J. Effects of metformin and other biguanides on oxidative phosphorylation in mitochondria. Biochem J 462, 475-487, doi:10.1042/BJ20140620 (2014).

21 Matsuzaki, S. & Humphries, K. M. Selective inhibition of deactivated mitochondrial complex I by biguanides. Biochemistry 54, 2011-2021, doi:10.1021/bi501473h (2015).

22 Luengo, A., Sullivan, L. B. & Heiden, M. G. Understanding the complex-I-ty of metformin action: limiting mitochondrial respiration to improve cancer therapy. BMC Biol 12, 82, doi:10.1186/s12915-014-0082-4 (2014).

23 Ludbrook, G. L., Visco, E. & Lam, A. M. Propofol: relation between brain concentrations, electroencephalogram, middle cerebral artery blood flow velocity, and cerebral oxygen extraction during induction of anesthesia. Anesthesiology 97, 1363-1370 (2002).

24 Vanlander, A. V. et al. Possible pathogenic mechanism of propofol infusion syndrome involves coenzyme q. Anesthesiology 122, 343-352, doi:10.1097/ALN.0000000000000484 (2015).

25 Krajcova, A., Waldauf, P., Andel, M. & Duska, F. Propofol infusion syndrome: a structured review of experimental studies and 153 published case reports. Crit Care 19, 398, doi:10.1186/s13054-015-1112-5 (2015).

26 Yang, N., Liang, Y., Yang, P., Yang, T. & Jiang, L. Propofol inhibits lung cancer cell viability and induces cell apoptosis by upregulating microRNA-486 expression. Braz J Med Biol Res 50, e5794, doi:10.1590/1414-431X20165794 (2017).

27 Konno, A. et al. Continuous monitoring of caspase-3 activation induced by propofol in developing mouse brain. Int J Dev Neurosci 51, 42-49, doi:10.1016/j.ijdevneu.2016.04.007 (2016).

28 Meng, C. et al. Propofol induces proliferation partially via downregulation of p53 protein and promotes migration via activation of the Nrf2 pathway in human breast cancer cell line MDA-MB-231. Oncol Rep 37, 841-848, doi:10.3892/or.2016.5332 (2017).

29 Cui, W. Y., Liu, Y., Zhu, Y. Q., Song, T. & Wang, Q. S. Propofol induces endoplasmic reticulum (ER) stress and apoptosis in lung cancer cell H460. Tumour Biol 35, 5213-5217, doi:10.1007/s13277-014-1677-7 (2014).

30 Kroemer, G., Dallaporta, B. & Resche-Rigon, M. The mitochondrial death/life regulator in apoptosis and necrosis. Annu Rev Physiol 60, 619-642, doi:10.1146/annurev.physiol.60.1.619 (1998).

31 Kajimoto, M. et al. Propofol compared with isoflurane inhibits mitochondrial metabolism in immature swine cerebral cortex. J Cereb Blood Flow Metab 34, 514-521, doi:10.1038/jcbfm.2013.229 (2014).

32 Forkink, M. et al. Complex I and complex III inhibition specifically increase cytosolic hydrogen peroxide levels without inducing oxidative stress in HEK293 cells. Redox Biol 6, 607-616, doi:10.1016/j.redox.2015.09.003 (2015).

33 Wong, H. S., Dighe, P. A., Mezera, V., Monternier, P. A. & Brand, M. D. Production of superoxide and hydrogen peroxide from specific mitochondrial sites under different bioenergetic conditions. J Biol Chem 292, 16804-16809, doi:10.1074/jbc.R117.789271 (2017).

34 Brand, M. D. Mitochondrial generation of superoxide and hydrogen peroxide as the source of mitochondrial redox signaling. Free Radic Biol Med 100, 14-31, doi:10.1016/j.freeradbiomed.2016.04.001 (2016).

35 Koshikawa, N., Hayashi, J., Nakagawara, A. & Takenaga, K. Reactive oxygen species-generating mitochondrial DNA mutation up-regulates hypoxia-inducible factor-1alpha gene transcription via phosphatidylinositol 3-kinase-Akt/protein kinase C/histone deacetylase pathway. J Biol Chem 284, 33185-33194, doi:10.1074/jbc.M109.054221 (2009).

36 Cohen, P. J. Effect of anesthetics on mitochondrial function. Anesthesiology 39, 153-164 (1973).

37 Ishikawa, K. & Hayashi, J. A novel function of mtDNA: its involvement in metastasis. Ann N Y Acad Sci 1201, 40-43, doi:10.1111/j.1749-6632.2010.05616.x (2010).

38 El-Mir, M. Y. et al. Dimethylbiguanide inhibits cell respiration via an indirect effect targeted on the respiratory chain complex I. J Biol Chem 275, 223-228 (2000).

39 Owen, M. R., Doran, E. & Halestrap, A. P. Evidence that metformin exerts its anti-diabetic effects through inhibition of complex 1 of the mitochondrial respiratory chain. Biochem J 348 Pt 3, 607-614 (2000).

40 Drahota, Z. et al. Biguanides inhibit complex I, II and IV of rat liver mitochondria and modify their functional properties. Physiol Res 63, 1-11 (2014).

41 Gottlieb, A., Duberstein, J. & Geller, A. Phenformin acidosis. N Engl J Med 267, 806-809, doi:10.1056/NEJM196210182671604 (1962).

42 Relman, A. S. Lactic acidosis and a possible new treatment. N Engl J Med 298, 564-565, doi:10.1056/NEJM197803092981009 (1978).

43 Finsterer, J. & Scorza, F. A. Effects of antiepileptic drugs on mitochondrial functions, morphology, kinetics, biogenesis, and survival. Epilepsy Res 136, 5-11, doi:10.1016/j.eplepsyres.2017.07.003 (2017).

44 Zhang, C. et al. Valproic Acid Promotes Human Glioma U87 Cells Apoptosis and Inhibits Glycogen Synthase Kinase-3beta Through ERK/Akt Signaling. Cell Physiol Biochem 39, 2173-2185, doi:10.1159/000447912 (2016).

45 Uppala, R. et al. Aspirin increases mitochondrial fatty acid oxidation. Biochem Biophys Res Commun 482, 346-351, doi:10.1016/j.bbrc.2016.11.066 (2017).

46 Moreno-Lastres, D. et al. Mitochondrial complex I plays an essential role in human respirasome assembly. Cell Metab 15, 324-335, doi:10.1016/j.cmet.2012.01.015 (2012).

47 Okamoto, A. et al. HIF-1-mediated suppression of mitochondria electron transport chain function confers resistance to lidocaine-induced cell death. Sci Rep 7, 3816, doi:10.1038/s41598-017-03980-7 (2017).

48 Yokota, M. et al. Generation of trans-mitochondrial mito-mice by the introduction of a pathogenic G13997A mtDNA from highly metastatic lung carcinoma cells. FEBS Lett 584, 3943-3948, doi:10.1016/j.febslet.2010.07.048 (2010).

49 Okamoto, A. et al. The antioxidant N-acetyl cysteine suppresses lidocaine‐ induced intracellular reactive oxygen species production and cell death in neuronal SH-SY5Y cells. BMC Anesthesiol 16, 104, doi:10.1186/s12871-016-0273-3 (2016).

50 Cheng, G. et al. Mitochondria-Targeted Analogues of Metformin Exhibit Enhanced Antiproliferative and Radiosensitizing Effects in Pancreatic Cancer Cells. Cancer Res 76, 3904-3915, doi:10.1158/0008-5472.CAN-15-2534 (2016).

51 Salabei, J. K., Gibb, A. A. & Hill, B. G. Comprehensive measurement of respiratory activity in permeabilized cells using extracellular flux analysis. Nat Protoc 9, 421-438, doi:10.1038/nprot.2014.018 (2014).

